# Single-nucleus transcriptional and chromatin accessibility analyses of maturing mouse Achilles tendon uncover the molecular landscape of tendon stem/progenitor cells

**DOI:** 10.1101/2024.10.24.619991

**Authors:** Hiroki Tsutsumi, Tomoki Chiba, Yuta Fujii, Takahide Matsushima, Tsuyoshi Kimura, Akinori Kanai, Akio Kishida, Yutaka Suzuki, Hiroshi Asahara

**Author notes:** These authors contributed equally to this work (Hiroki Tsutsumi, Tomoki Chiba).

## Abstract

Tendons and ligaments are crucial connective tissues linking bones and muscles, yet achieving full functional recovery after injury remains challenging. We investigated the characteristics of tendon stem/progenitor cells (TSPCs) by focusing on the declining tendon repair capacity with growth. Using single-cell RNA sequencing on Achilles tendon cells from 2-and 6-week-old mice, we identified *Cd55* and *Cd248* as novel surface antigen markers for TSPCs. Combining single-cell RNA sequencing with single-nucleus RNA and ATAC sequencing analyses revealed that *Cd55* and *Cd248* positive fractions in tendon tissue represent TSPCs, as confirmed by their expression of established TSPC markers, with this population decreasing at 6 weeks. We also identified candidate upstream transcription factors regulating these fractions. Functional analyses of isolated CD55/CD248 positive cells demonstrated high clonogenic potential and tendon differentiation capacity, forming functional tendon-like tissue *in vitro*. This study establishes CD55 and CD248 as novel TSPC surface antigens, potentially advancing tendon regenerative medicine and contributing to the development of new treatment strategies for tendon and ligament injuries.

## Introduction

Tendons are connective tissues involved in the contractile movement of muscle to bone. Tendons are rich in extracellular matrix components, such as type 1 collagen and proteoglycans, which both have elastic and viscous properties to withstand overload (Andarawis-Puri et al., 2015; Asahara et al., 2017; Connizzo et al., 2013).

Tendons have a hierarchical structure. Collagen fibrils aggregate to form fibers, which then bundle into fascicles. Each fascicle is surrounded by a thin connective tissue layer called the endotenon. Fascicles are further enclosed by the epitenon, and in some cases, an additional outer layer called the paratenon. The number of fascicles and the presence of a paratenon can vary among different tendons and species (Jozsa et al., 1991; Walia et al., 2019).

Tendon and ligament injuries potentially account for the majority of musculoskeletal disorders and can lead to arthritis and spondylitis. (Gracey et al., 2020). The most popular treatment approaches are surgery and conservation (Steinmann et al., 2020). However, full functional recovery is often not achieved due to scarring and fibrosis in the injured area (Guilak et al., 2014; Thomopoulos S et al., 2015). Furthermore, the risk of postoperative rupture is high (Leong et al., 2020; Loiacono et al., 2019). In this context, tendon regeneration focused on tendon stem/progenitor cells (TSPCs) is attracting attention (Leong et al., 2016). Tendons have few cellular components, including tendon cells (tenocytes) and TSPCs (Tang et al., 2016; Bi et al., 2007). Tenocytes are responsible for tendon homeostasis, while TSPCs self-renew and differentiate into tenocytes. Thus, investigating TSPCs biology is important for understanding tendon regeneration and homeostasis.

Murine TSPCs were first reported in 2007 by Bi Y et al. (Bi et al., 2007). They demonstrated that TSPCs possess self-renewal capacity, colony-forming ability, and multi-differentiation potential *in vitro*. Furthermore, murine TSPCs were reported to simultaneously express stem cell markers such as *Cd44* and *Stem cells antigen-1* (*Ly6a*), as well as tendon-related genes like *Scleraxis* (*Scx)* and Collagen type I alpha 1 chain (*Col1a1*), and were identified as a subset of tendon cells within the tendon fascicle.

Subsequent research on murine TSPC localization has yielded diverse perspectives. Harvey et al. reported that *Tppp3*/*Pdgfra*-positive cells in the epitenon are induced upon tendon injury and contribute to repair (Harvey et al., 2019). Additionally, Yin et al. and Tempfer et al. demonstrated the presence of TSPCs around blood vessels (Yin et al., 2016; Tempfer et al., 2009).

Markers used to characterize murine TSPCs include CD73, CD105, and CD90, which are known criteria for mesenchymal stem cells (MSCs) (Lui et al., 2015). Additionally, TSPCs have been reported to express *POU class 5 homeobox 1* (*Oct-4*), *Nanog*, *Nucleostemin*, stage-specific embryonic anitigen-4 *(SSEA-4)*, *c-Myc*, *SRY-box transcription factor (Sox*2), *Fucosyltransferase 4* (*Fut-4*), and other genes (Zhang et al., 2016). Expression of CD146 and CD44 has been confirmed as well (Ruzzini et al., 2014). The *Tppp3*/*Pdgfra*-positive TSPCs reported by Harvey et al. highly express *Cd34*, which is generally considered to have low expression in MSCs (Harvey et al., 2019; Tachibana, et al., 2022). *Cd34* is known to be highly expressed in mouse embryonic limb buds at E14.5 compared to E11.5 (Havis et al., 2014), suggesting that *Cd34* positive cells might reflect progenitor cells that constitute the limb bud, including tendons. However, these markers are not specific to TSPCs, thus, a definitive method to distinguish TSPCs from mature tendons *in vivo* has not been established yet (Li et al., 2019; Wang et al., 2008; Chen et al., 2012; Cho et al., 2018; Fang et al., 2022).

Regarding tendon regenerative capacity, Howell et al. reported an interesting observation. Tendons in juvenile mice can regenerate functional tissue after injury, but this ability is lost in mature mice, resulting in scar tissue formation (Howell et al., 2017). This finding suggests the possibility of abundant TSPCs in juvenile mouse tendons.

Given this background, we hypothesized that evaluating tendon tissue heterogeneity at the single-cell level using juvenile mouse tendons would enable a more detailed characterization of TSPCs. This approach is expected to advance the identification and characterization of TSPCs, which has been challenging with conventional methods.

Therefore, we performed scRNA-seq using cells collected from 2-and 6-week-old mouse Achilles tendons and investigated clusters that co-express known TSPCs markers, such as *Tppp3*, *Pdgfra*, and *Ly6a*. Then, we analyzed the expression of surface antigens in these clusters and identified *Cd55* and *Cd248* as novel candidate surface antigens for murine TSPCs.

Furthermore, snATAC-seq and snRNA-seq were performed simultaneously using cells collected from Achilles tendons to evaluate the validity of *Cd55* and *Cd248* as surface antigens for TSPCs and to identify the landscape of transcription factors (TFs) involved in tendon maturation. We sorted mouse Achilles tendon cells based on CD55 and CD248 and confirmed their phenotypes, demonstrating high clonogenicity and highly efficient induction into tendon cells. These results suggested that CD55 and CD248 are novel surface antigens of murine TSPCs and may be useful for understanding the process of tendon maturation and for applications in cell therapy.

## Material and Methods

### Single cell isolation

Mice were euthanized by exposure to CO_2_ followed by cervical dislocation. To remove Achilles tendons, a longitudinal incision through the skin was made down the midline of the posterior aspect of the lower limb. A sharp transverse incision was made just distal to the myotendinous junction and again just proximal to the enthesis, and the Achilles tendon was carefully removed. The procedure was performed bilaterally. All animals were purchased from Sankyo Lab Service (Tokyo, Japan). Achilles tendons were digested to obtain a single-cell suspension. In total, 40 or 60 (2-week and 6-week, respectively) mouse Achilles tendons were processed together as a single sample. After washing with 1× phosphate-buffered saline (PBS) several times and cutting 1-mm-wide sections using scissors, tendons were digested for 1 h at 37°C with continuous shaking at 1,200 rpm and 37°C in a dissociation solution consisting of 30 mg/mL collagenase (Wako, Osaka, Japan). After digestion, the single-cell suspension was filtered for debris using a 40 μm cell strainer and washed twice with 1× PBS. The samples were centrifuged for 15 min at 1,500 rpm and 4°C. Dead cells were removed using a Dead Cell Removal Kit (Miltenyi Biotec, Bergisch Gladbach, Germany). Cell viability was 76% (2-week) and 73% (6-week), respectively. The experiment involving mice was approved by Animal Experimental Committee of the Institute of Science Tokyo.

### scRNA-seq library construction and sequencing

Cells isolated from Achilles tendons of both 2-week-old and 6-week-old mice were processed separately. Cells were resuspended in PBS with 1% fetal bovine serum (FBS) at a concentration of 10,000 cells per μL. Then, 10,000 cells per sample were loaded on a Chromium Controller (10x Genomics, Pleasanton, CA, USA) for single cell capture.

Libraries were prepared using Single Cell 3′ Library & Gel Bead Kit v3 (10x Genomics) following the manufacturer’s instructions. A single-cell emulsion (Gel Bead-In-EMulsions, GEMs) was created by making barcoded cDNA unique to each individual emulsion. A recovery agent was added to break GEM and cDNA was then amplified. A library was produced via end repair, dA-tailing, adapter ligation, post-ligation cleanup with SPRIselect, and sample index PCR. The quality and concentration of the amplified cDNA were evaluated using the Bioanalyzer (Agilent 2100) on a High Sensitivity DNA chip (Agilent, #5065-4401; Santa Clara, CA, USA). Sequencing was performed using the HiSeq X system (Illumina, San Diego, CA, USA) to generate 28/90 bp paired-end reads.

### Cell dissociation, nuclei isolation, and snRNA-seq and snATAC-seq library construction and sequencing

After obtaining cells from the mouse Achilles tendons of 2-week-old and 6-week-old mice (processed separately for each age group), nuclei were isolated following the Chromium Next GEM Single Cell Multiome ATAC + Gene Expression Reagent Bundle (10XGenomics) and all of the buffers were made according to the manufacturer’s instructions. Briefly, after centrifugation, the supernatant was discarded, and the cells were re-suspended in 1 mL of 1% FBS in PBS. Approximately 500,000 cells were transferred to a new tube for further lysis. To remove the supernatant, cells were centrifuged again for 5 min at 300 rcf and 4°C and resuspended in 100 mL of chilled lysis buffer. Cells were lysed for 9 min on ice and 1 mL of wash buffer was added to stop the reaction. Then, the suspension was filtered through a 40 μm cell strainer, cells were centrifuged and resuspended in 66.2 mL of chilled Diluted Nuclei Buffer aiming to target 7000 nuclei. The final concentration of nuclei was determined, followed by transposition, GEM generation, barcoding, and library construction according to the Chromium Next GEM Single Cell Multiome ATAC + Gene Expression (10XGenomics). Libraries were sequenced with the parameters recommended by the manufacturer, using the NovaSeq 6000 (Illumina) to generate 28/90 bp paired-end reads for gene expression and 50/49 bp paired-end reads for ATAC sequencing.

### scRNA-seq and snRNA-seq analyses

Sequencing reads were processed with the Cellranger_arc (10X Genomics, v2.0.0) using the mouse reference mm10. We performed separate analyses for scRNA-seq data (collected from both 2-week and 6-week samples) and snRNA-seq data (collected from both 2-week and 6-week samples) to enable comprehensive characterization of transcriptional landscapes. From the gene expression matrix, downstream analyses were carried out using R. Quality control, filtering, data clustering and visualization, and a differential expression analysis were carried out using Seurat (Butler et al., 2018). For each dataset, cells with unique feature counts over 2,500 or less than 200 and >5% mitochondrial counts were filtered. Then, heterotypic doublets (assuming 5% of barcodes represent doublets) were removed using DoubletFinder (McGinnis et al., 2019).

Unsupervised shared nearest neighbor (SNN) clustering was performed with varying resolution and the results were visualized using uniform manifold approximation and projection (UMAP) (Becht et al., 2019). Differentially expressed genes (DEGs) among each cell cluster were determined using the FindAllMarkers function in Seurat. The criteria of DEGs as a marker gene for the cluster was logFC > 0.25, adjusted p < 0.05, expression in >25% of cells. We then annotate each cluster based on DEGs (supplementary table1-15). To ensure consistency between our different sequencing approaches, we compared cell clusters identified in the scRNA-seq analysis with those from the snRNA-seq analysis (Supplemental Figure 5).

### Pseudotime analysis

Monocle3 (Trapnell et al., 2014) was used to convert the snRNA-seq dataset into a cell dataset object (CDS), preprocess data, correct for batch effects, embed with dimensional reduction, and perform pseudotemporal ordering. Cicero (Pliner et al., 2018) was used to generate pseudo-temporal trajectories for the snATAC-seq dataset.

### Cell-cell communication analysis

Intercellular communication networks were quantitatively inferred and analyzed using scRNA-seq data. The R package CellChat (Jin et al., 2021) was used to visualize the interactions among different cell groups. Two hundred twenty-nine signaling pathway families were grouped as a library to analyze cell–cell communication.

### GO analysis

The R package ClusterProfiler (Yu et al., 2012) was used to perform a gene enrichment analysis. The p-values were corrected by the Benjamini & Hochberg method.

### Motif enrichment analysis (ChiP seeker)

Genomic regions containing snATAC-seq peaks were annotated using ChIPSeeker (Yu et al., 2015) and the UCSC database on mouse (mm10).

### snATAC-seq analysis

The Cell Ranger ATAC pipeline (1.2.0) (Satpathy et al., 2019) was used to preprocess the data resulting from sequencing. First, Tn5 sites were mapped to the mouse reference transcriptome mm10, and duplicate reads and background cells were removed. This returned barcoded fragment files, which were loaded into Signac (Stuart et al., 2021) for downstream analyses using the standard Signac/Seurat pipeline. Macs2 (Zhang et al., 2008) was run on the fragment files to call peaks using the Signac ‘CallPeaks’ function.

Fragments were mapped to the Macs2 called peaks and assigned to cells using the Signac ‘FeatureMatrix’ function. Nucleosome signal strength and transcription start site (TSS) enrichment for each cell were calculated using the Signac ‘NucleosomeSignal’ and ‘TSSEnrichment’ functions, respectively. Outliers in the QC metric categories were removed as per Signac’s standard processing guidelines. Latent semantic indexing (LSI), a form of dimensional reduction, was performed using the Signac ‘RunTFIDF’ and ‘RunSVD’ functions. The UMAP hyperparameters were varied to produce consistent object shapes (using R). Once hyperparameters were chosen, the Signac/Seurat’s ‘RunUMAP’ function was run on the LSI dimensions chosen earlier for UMAP embedding. The Signac/Seurat ‘FindNeighbors’ function was run using the same LSI dimensions used for UMAP to compute the nearest neighbor graph. Signac/Seurat ‘FindClusters’ was then run at varying resolutions. A gene activity matrix was constructed by counting ATAC peaks within the gene body and 2 kb upstream of the transcriptional start site for protein-coding genes annotated in the Ensembl database. The gene activity matrix was log-normalized prior to label transfer with the aggregated snRNA-seq Seurat object using a canonical correlation analysis. The aggregated snATAC-seq object was filtered using a 97% confidence threshold for cell-type assignment following label transfer to remove heterotypic doublets. The filtered snATAC-seq object was reprocessed with TFIDF, SVD, and batch effect correction followed by clustering and annotation based on lineage-specific gene activity. Differential chromatin accessibility between cell types was assessed with the Signac FindMarkers function for peaks detected in at least 20% of cells using a likelihood ratio test and a log-fold-change threshold of 0.25. Bonferroni-adjusted p-values were used to determine significance at an FDR < 0.05.

### Integration of snRNA-seq and snATAC-seq data

snRNA-seq and snATAC-seq data were integrated using the cluster-label transfer procedure as implemented in Signac and Seurat. Each snRNA-seq sample. as clustered individually and its cluster labels were projected onto the matching, individually clustered snATAC-seq sample or vice versa. Anchors were identified for condition-matched snRNA-and snATAC-seq samples using the FindTransferAnchors function and a canonical correlation analysis (CCA) was performed using the snRNA expression values and the snATAC-imputed gene expression values. The anchors were used to transfer cluster-label identifiers between the two data types using the TransferData function. Each cell in the query was assigned the cluster label with the highest prediction score, and label transfer was considered successful for query cells with prediction scores above 0.3.

### SCENIC analysis

To identify TFs and characterize cell states, a cis-regulatory analysis was performed using the R package SCENIC (Lake et al., 2018), which infers gene regulatory networks based on co-expression and DNA motif analyses. The network activity was then analyzed for each cell to identify recurrent cellular states. In short, TFs were identified using GENIE3 and compiled into modules (regulons), which were subsequently subjected to cis-regulatory motif analysis using RcisTarget with two gene-motif rankings: 10 kb around the TSS and 500 bp upstream. Regulon activity in every cell was then scored using AUCell.

### Immunohistochemistry

Immunohistochemistry was performed as previously described (Tsutsumi et al., 2022). Briefly, tissue samples were fixed in 4% paraformaldehyde overnight at 4°C and embedded in paraffin. Sections of 10 μm in thickness were deparaffinized and activated with citric acid buffer (10 mM sodium citrate, 1 mM EDTA; pH 6.0) at 80 °C for 60 min in a decloaking chamber NxGen (BIOCARE Medical, CA, USA) or with 0.1% trypsin/PBS at 37 °C for 30 min. After they were blocked with Blocking One solution (Nacalai tesque, Kyoto, Japan) for 60 min, they were incubated with rabbit anti-CD55 antibody (1:100; A13918, ABclonal, Woburn, MA, USA) or rat anti-CD248 (1:100; MAB7535, R&D Systems, Minneapolis, MN, USA) overnight at 4 °C. Following this step, they were incubated with Alexa Fluor plus 488 goat anti-rabbit antibody (1:1000; A32731, Thermo Fisher Scientific, Waltham, MA, USA) or Alexa Fluor plus 488 donkey anti-rat antibody (1:1000; A21208, Thermo Fisher Scientific) for 60 min. Hoechst 33342 (100ng/ml, Thermo Fisher Scientific) was added during this incubation. The sections were then mounted with ProLong Glass Antifade Mountant (P36980, Thermo Fisher Scientific).

### Fluorescence-activated cell sorting (FACS)

Cells harvested from the Achilles tendon were incubated for 60 min on ice with APC-CD55 (#131812; BioLegend, San Diego, CA, USA) and a primary antibody against CD248 (#LS-B2712, LSBio) using a FACS buffer (PBS containing 1 [v/v%] FBS). After washing, cells were reacted with fluorescent-conjugated rabbit secondary antibody. Cell sorting was performed using MoFlo XDP FACS (Beckman Coulter, Brea, CA, USA). DAPI was used as the live/dead discriminator to the gate. CD55/CD248 dual positive and negative cell cluster were sorted.

### Primary cell culture

Primary TSPCs were isolated from the Achilles tendons of 2-week-old mice with collagenase digestion described above. Single-cell suspensions were cultured in the culture medium (MEMα + 20% FBS + 1% penicillin-streptomycin + 1 [v/v%] 100× non-essential amino acid solution [NEAA; Gibco], 1 [v/v%] 100× GlutaMAX [Gibco]). At 80%–90% confluence, cells were trypsinized, centrifuged, resuspended in culture medium as passage 1 cells, and incubated in 5% CO_2_ at 37°C, with fresh medium every 2–3 days.

### Colony formation assay

For the colony formation assay, single-cell suspensions of TSPCs (1000 cells/well) were seeded and incubated in six-well plates for 12–14 days in the growth medium and fixed with 4% paraformaldehyde (PFA) (Sigma-Aldrich, St. Louis, MO, USA). Then, 0.1% crystal violet solution (Wako) was used to stain the cells. Colonies of >30–50 cells were defined as a single colony unit, and the number of clusters was counted using the ImageJ package ColonyArea (Guzmán et al., 2014)

### RT-PCR

Total RNA was purified using TRIzol reagent (Invitrogen, Grand Island, NY, USA). Reverse transcription of mRNA was carried out using the PrimeScript RT Reagent Kit (Takara, Shiga, Japan). Quantitative PCR (qRT-PCR) was performed on cDNA with the Thunderbird SYBR mix (Toyobo, Osaka, Japan). B2m was used as a reference gene, and relative gene expression levels were calculated through the ΔCT method.

### Tenocytes, cartilage, and osteocyte differentiation

For the differentiation experiment, TSPCs were cultured in 6-well plates (50,000 cells/well). Osteogenic, chondrogenic, and tenogenic differentiation were induced using a corresponding differentiation medium. The osteogenic differentiation medium contained a culture medium supplemented with 10 nM dexamethasone (Sigma-Aldrich), 5 mM β-glycerophosphate (APEXBIO), and 0.05 mM L-ascorbic acid 2-phosphate (Sigma-Aldrich). The chondrogenic differentiation medium contained a culture medium supplemented with 100 nM dexamethasone and 10 ng/mL BMP2 (Sigma-Aldrich). The tenogenic differentiation medium contained a culture medium supplemented with 10 ng/mL TGF-β1 (Peprotech, Rocky Hill, NJ, USA), 10 ng/mL GDF-5 (R&D Systems, Minneapolis, MN, USA), and 0.05 mM L-ascorbic acid 2-phosphate. After a 2 week induction period, cells were harvested for RT-PCR.

### Bio-cultured tendon (Bio-tendon)

Generation of Bio-tendon has reported previously (Kataoka et al., 2020; Tsutsumi et al., 2022) Briefly, sorted cells were embedded in a 3D-culture cocktail, which is consistent with 2 mg/mL Cellmatrix (Type I-A, Nitta Gelatin Inc., Osaka, Japan), pro-survival cocktail (final concentrations: 100 nM Bcl-Xl BH4 4-23 (Merck Millipore, Burlington, MA, USA), 100 μM carbobenzoxy-valyl-alanyl-aspartyl-[*O*-methyl]-fluoromethylketone (Z-VAD-FMK) (Promega, Madison, WI, USA)) in culture medium. The 3D chamber was coated with 1% gelatin and incubated at 37°C and 5% CO_2_ for 30 min. After washing with 1× PBS 3 times, 1.0 × 10^6^ cells were transferred to a 3D-culture cocktail mixture and incubated at 37°C and 5% CO_2_ for 60 min for gelation. Following gelation, tenogenic differentiation medium was added to the chamber. Following 24 h of incubation, the chamber was placed within a cell stretching system (Shellpa Pro, Menicon Life Science, Aichi, Japan). Cyclic mechanical stretch was applied for 1 week, with gradual increase in stretch load: 1% on day 1, 2% on day 2, 3% on day 3, 4% on day 4, and 5% from days 5 to 7. The stretching cycle was programed at 0.25 Hz for 18 h/day, followed by resting for 6 h/day at 37°C and 5% CO_2_. Daily medium changes were also required.

### Decellularization by HHP and chemical treatment

The cultured bio-tendon was placed into a plastic pack filled with saline and sealed to prevent implosion and leakage during the procedure. The pack was then pressurized at 1,000 MPa at 30°C for 10 min using an HHP machine (Dr. CHEF; Kobe Steel, Hyogo, Japan). After pressurization, the bio-tendon was washed thrice with 30% ethanol (EtOH) by continuous shaking for 5 min at each step. Finally, 1-ethyl-3-(3-dimethylaminopropyl) carbodiimide (EDC)/*N*-hydroxysuccinimide (NHS)-based cross-linking was performed by adding 70 mM EDC (Wako) and 70 mM NHS (Wako) with 30% EtOH for 24 h at 4°C. The processed bio-tendon was incubated in 1× PBS at 4°C until the experiment.

### Electron microscopy

Bio-tendon tissues were dissected and fixed overnight in 2.5% glutaraldehyde in 0.1 M phosphate buffer (PB). For transmission electron microscopy (TEM), the specimens (*n* = 3) were rinsed with 0.1 M PB, post-fixed in 1% osmium buffered with 0.1 M PB for 2 h, and dehydrated in a graded ethanol series. The specimens were then embedded in Epon 812, sectioned into ultrathin sections (70 nm), mounted on copper grids, and double-stained with uranyl acetate and lead citrate. TEM observation was conducted JEM-1400Flash (JEOL, Tokyo, Japan). For scanning electron microscopy (SEM), the specimens (*n* = 2) were dried in a critical-point dryer (HCP-2; Hitachi, Tokyo, Japan) with liquid CO_2_ and coated with platinum. The specimens were subsequently observed under SEM (JSM-7900F/JED-2300; JEOL). Fiber orientation within the microscopic sections was analyze using the OrientationJ image processing tool, a plugin for ImageJ.

### Stretch test

The mechanical properties were measured using a creep meter (RE-3305S; Yamaden, Tokyo, Japan). After measuring the initial length (mm), diameter (mm), and thickness (mm) using a micrometer, the samples were fixed with two grips, which were pulled at a constant speed of 0.05 mm/s until failure, and the tensile strength (N) and failure strain (mm) were measured. The cross-sectional area (mm^2^) was calculated using the initial diameter and thickness. The stiffness was manually determined from the slope in the linear region of the failure-stress curve.

The tensile strength (MPa), failure strain (%), stiffness, and Young’s modulus were calculated using the following formulas:

Tensile strength (MPa) = tensile strength (N)/cross-sectional area (mm^2^). Failure strain (%) = failure strain (mm)/initial length (mm).

Stiffness = Stress (N)/Strain (mm)

Young’s modulus = stiffness × initial length (mm)/cross-sectional area (mm^2^)

## Statistical analysis

All values are presented as means ± SEM. Statistically significant differences were assessed using unpaired two-tailed Student’s *t*-tests and one-way analysis of variance (ANOVA) with Tukey’s post hoc tests. Statistical significance was set at p < 0.05.

## Results

### Identification of Novel Surface Antigens for Tendon Stem/Progenitor Cells (TSPCs)

In mice, the healing capacity for tendon injuries is high up to 2 weeks of age, but decreases as the musculoskeletal system matures at 6 weeks (Howell et al., 2017). To investigate whether this phenomenon is due to changes in the cellular population of tendon tissue, including fluctuations in progenitor cells, we performed single-cell RNA sequencing on mouse Achilles tendons at these two time points.

Achilles tendons were harvested from mice at 2 and 6 weeks of age. Following collagenase digestion and dead cell removal, we employed massively parallel, droplet-enabled scRNA-seq analysis (10X Genomics Chromium). Data processing was conducted using the Cell Ranger pipeline (10X Genomics). We analyzed 10,314 cells from 2-week-old mice (median 3,167 genes/cell and 41,611 mean reads/cell) and 6,513 cells from 6-week-old mice (median 732 genes/cell and 60,386 mean reads/cell). After doublet removal using Doubletfinder, we merged the datasets and performed unbiased clustering using Seurat, identifying a total of 15 clusters (Fig. 1A). The top differentially expressed genes (DEGs) for each cluster, based on log2 fold change and statistical significance, are summarized in Fig. 1B.

**Figure 1.**
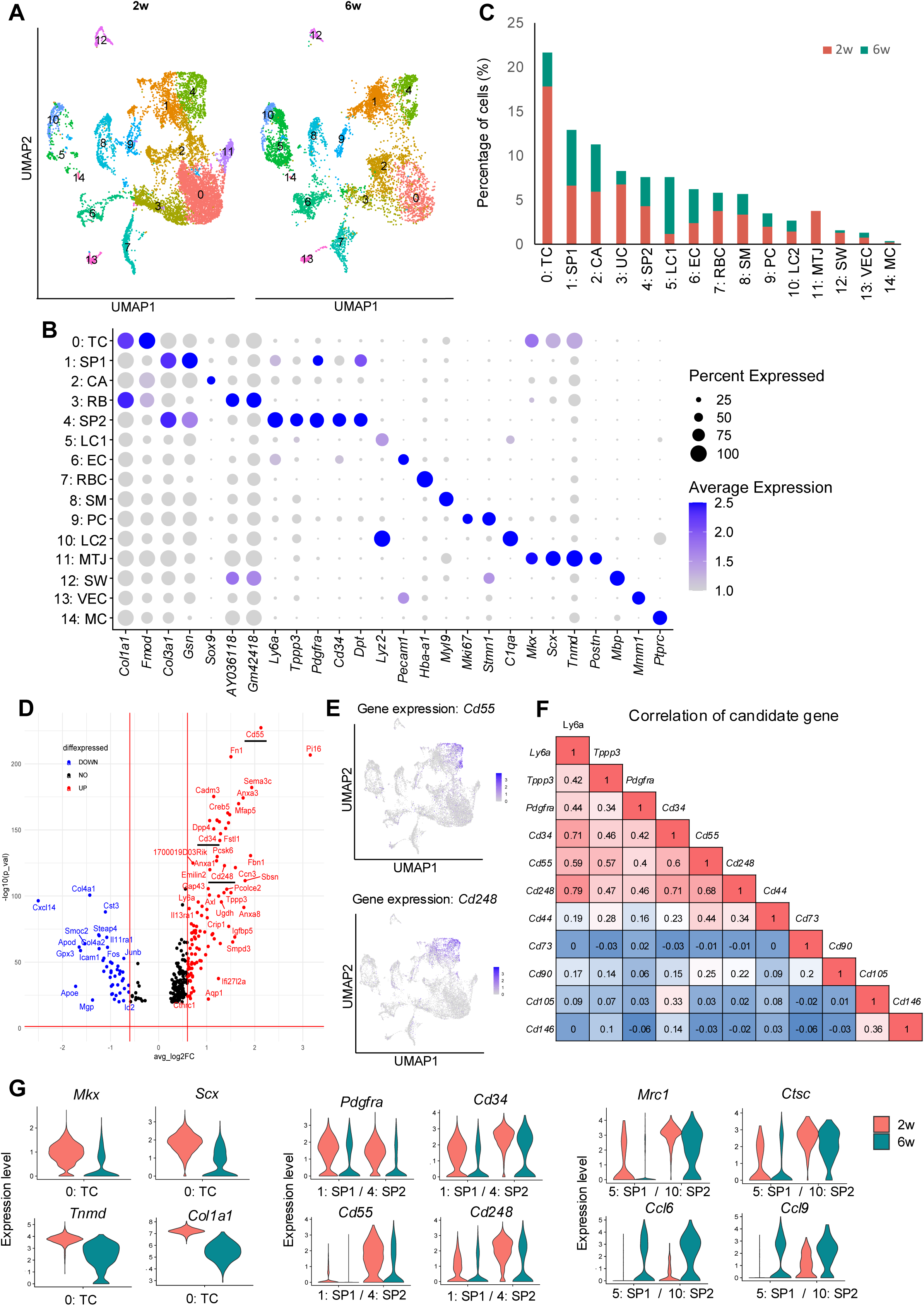
scRNA-seq of tendon cells from 2-week-old and 6-week-old mice and the identification of surface markers of TSPC. (A) Integrated uniform manifold approximation and projection (UMAP) scRNA-seq clustering of cells harvested from 2-week-old and 6-week-old mouse Achilles tendons. (B) Dot plot of average gene expression levels of the indicated genes in each scRNA-seq cluster. The size of the dot reflects the percentage of cells in the cluster that express each gene. TC, tenocyte; SP1, tendon stem/progenitor cell_1; CA, cartilage; RB, ribosomal RNA; SP2, tendon stem/progenitor cell_2; LC1, lymphocyte_1; EC, endothelial cell; RBC, red blood cell; SM, smooth muscle cell; PC, proliferating cell; LC2, lymohocytes_2; MTJ, myotendinous junction cell; SW, Schwann cell; VEC, vascular endothelial cell; MC, macrophage. (C) Proportions of cells in clusters identified from scRNA-seq. Clusters are colored according to cluster type. (D) Volcano plot of gene expression in the SP2 cluster and the identification of candidate TSPC marker genes (red under line). (E) Feature plot of *Cd55* and *Cd248* expression. (F) Correlation of gene expression of TSPC candidate genes in 2-week data. (G) Violin plots presenting the gene expression changes for a selection of differentially expressed genes.

We identified two clusters (0 and 11) with high expression of *Mkx* and *Scx*, transcription factors involved in tendon development and growth. Cluster 0 showed higher expression of ECM-related genes such as *Col1a1* and *Fmod* compared to cluster 11, suggesting these were tenocytes (TC). Cluster 11, with high *Periostin* (*Postn)* expression (Jacobson, et al., 2020), was classified as myotendinous junction (MTJ). Other clusters were characterized as follows: cartilage (CA, high *SRY-box transcription factor 9* (*Sox9*) expression), lymphocytes (LC1, LC2, high *Lysozyme 2* (*Lyz2*) and *Complement component 1, q subcomponent, alpha polypeptide* (*C1qa*)), endothelial cells (EC, high *Platelet and endothelial cell adhesion molecule 1* (*Pecam1*)), red blood cells (RBC, high *Hemoglobin alpha, adult chain 1* (*Hba-a1*)), smooth muscle cells (SM, high *Myosin light chain 9* (*Myl9*)), proliferating cells (PC, high *Mki67* and *Stathmin 1* (*Stmn1*)), Schwann cells (SW, high *Myelin basic protein* (*Mbp*)), vascular endothelial cells (VEC, high *Multimerin 1* (*Mmrn1*)), and macrophages (MC, high *Protein tyrosine phosphatase receptor type C* (*Ptprc*)). In cluster3, *Col1a1* and *Fmod* were expressed; however, *Gm42418* and *AY036118* were highly expressed. These long non-coding RNAs are related to RN45s and therefore cluster3 represents ribosomal contamination (Isola et al., 2024). It is also speculated that the cartilage cluster (CA) reflects enthesis, with expression of *Scx* as well as *Sox9* (Zhan et al., 2023). DEG s of each cluster was summarized in supplementary table1-15.

To identify progenitor populations within these clusters, we analyzed expression patterns of previously reported markers *Tppp3* and *Pdgfra* (Harvey et al., 2019; Tachibana, et al., 2022), along with the known stem/progenitor cell marker *Ly6a* (Holmes et al., 2007; Sung et al., 2008; Hittinger et al., 2013; Sidney et al., 2014; Fang et al., 2022) (Supplemental Figure 1B). We identified subclusters within clusters 1 and 4 showing high expression of these genes, which we defined as SP1 and SP2. SP2 exhibited the highest expression of these genes, suggesting it had the strongest progenitor characteristics. It has been reported (Harvey et al., 2019) that the *Tppp3*-positive population is localized to peritenon, and SP clusters might reflect peritenon as well.

SP2 also showed strong expression of genes associated with early tendon development in mouse embryos, as identified by RNA-seq of mouse limb tendon cells at E14.5 (Havis et al., 2014) and the Eurexpress mouse embryo transcriptome atlas database (Diez-Roux et al., 2011) (Supplemental Figure 2A). Gene Ontology analysis revealed upregulation of tendon development-related pathways in SP2, including TGFβ production and collagen-containing extracellular matrix (Supplemental Figure 2B). Evaluation of cell numbers in each cluster showed a decrease in SP2, TC, and MTJ at 6 weeks (Fig. 1C, Supplemental Figure 1A).

We then examined DEGs in SP2 compared to SP1 (Fig. 1D). Whole list of DEGs comparing SP2 and SP1 was summarized in supplementary table16. Among these, we identified *Cd34*, *Cd55*, and *Cd248* as genes encoding surface antigens. *Cd34* is known to be highly expressed in mouse embryonic limb bud E14.5 compared to E11.5 (Havis et al., 2014), suggesting that *Cd34* positive cluster might reflect a progenitor cell that consist of the limb bud, including tendons. However, *Cd55* and *Cd248* have not been discussed with this context. Expression of *Cd55* and *Cd248* was localized to areas corresponding to SP1 and SP2 (Fig. 1E). We evaluated the correlation coefficients between the expression of these genes and previously suggested TSPC markers (*Cd73*, *Cd90*, *Cd105*, *Cd44*, *Cd146*) as well as *Tppp3*, *Pdgfra*, and *Ly6a* at 2 weeks of age. *Cd34*, *Cd55*, and *Cd248* showed high correlation, suggesting these genes as new candidate surface antigens for TSPCs (Fig. 1F).

Comparing gene expression between 2 and 6 weeks, we observed a decrease in *Mkx* and *Scx* expression, consistent with previous reports (Grinstein et al., 2019). Furthermore, the expression of our candidate tendon progenitor cell markers *Cd34*, *Cd55*, and *Cd248* all decreased at 6 weeks. Inflammatory cells, including macrophages, increased at 6 weeks (Fig. 1G). Gene expression analysis revealed a shift from M2 macrophage-related genes (*Mannose receptor C-type 1* (*Mrc1*) and *Cathepsin C* (*Ctsc*)) at 2 weeks to M1 macrophage-related genes (*Chemokine (C-C motif) ligand 6* (*Ccl6*) and *Chemokine (C-C motif) ligand 9* (*Ccl9*)) at 6 weeks. Comparative analysis between 2-week and 6-week tendons was summarized as a heatmap in supplemental Figure 3.

Analysis of cell-cell communication changes in postnatal tendon and surrounding tissues (Lui et al., 2019) using CellChat (Jin et al., 2021) showed a significant decrease in overall interaction at 6 weeks compared to 2 weeks. These results demonstrate that the cellular profile of tendon tissue changes dramatically at 6 weeks, with decreased expression of stem progenitor markers and reduced interaction with surrounding tissues.

### Single Nucleus RNA + ATAC Analysis of 2-Week-Old Mouse Achilles Tendon Cells

Gene regulation is mediated by the binding of transcription factors to cis-regulatory elements proximal to the gene. Consequently, epigenetic changes such as chromatin accessibility play a crucial role in gene expression (Miller et al., 2013). Moreover, transcription factors are typically expressed at low levels, potentially leading to false negatives in scRNA-seq due to detection limits. Chromatin accessibility changes often precede gene expression changes, potentially allowing us to predict transcriptional changes (Ranzoni et al., 2020). Therefore, we performed simultaneous ATAC-seq and RNA-seq at the single-cell level to identify gene expression changes and their associated epigenetic alterations, aiming to elucidate the post-natal growth mechanisms of tendon tissue and evaluate the validity of *Cd55* and *Cd248* as markers.

We extracted nuclei from cells isolated from 2-week-old mouse Achilles tendons and conducted droplet-enabled multi-omics analysis (10X Genomics Chromium), performing simultaneous snATAC-seq and snRNA-seq as a reference. We first analyzed the snATAC-seq data. After mapping reads to the genome and calling peaks, we annotated the location of peaks in terms of genomic features. The peaks were associated with promoters, introns, exons, and intergenic regions.

We evaluated 6,571 cells (median high-quality fragments per cell: 17,408) and clustered them using latent semantic indexing and Uniform Manifold Approximation and Projection (UMAP) with the R package Signac (Stuart et al., 2021) (Figure 2A). Given the limited knowledge of cell-specific chromatin accessibility, we assessed gene activity by computing counts per cell within the gene body and promoter of protein-coding genes. Using this gene activity data, we performed unsupervised clustering and identified 17 clusters (ground-truth annotation). Clusters expressing tendon-related genes such as *Mkx* and *Scx* were identified as clusters 1 and 6, defined as A1: TC1 and A6: TC2, respectively. Clusters 0, 14, and 15 were identified as expressing stem progenitor markers including *Cd34*, *Ly6a*, *Pdgfra*, *Cd55*, and *Cd248*, and were defined as A2-0: SP1, A2-14: SP2, and A2-15: SP3, respectively. The characteristic gene activities and annotations for each cluster are summarized in Fig. 2B.

**Figure 2.**
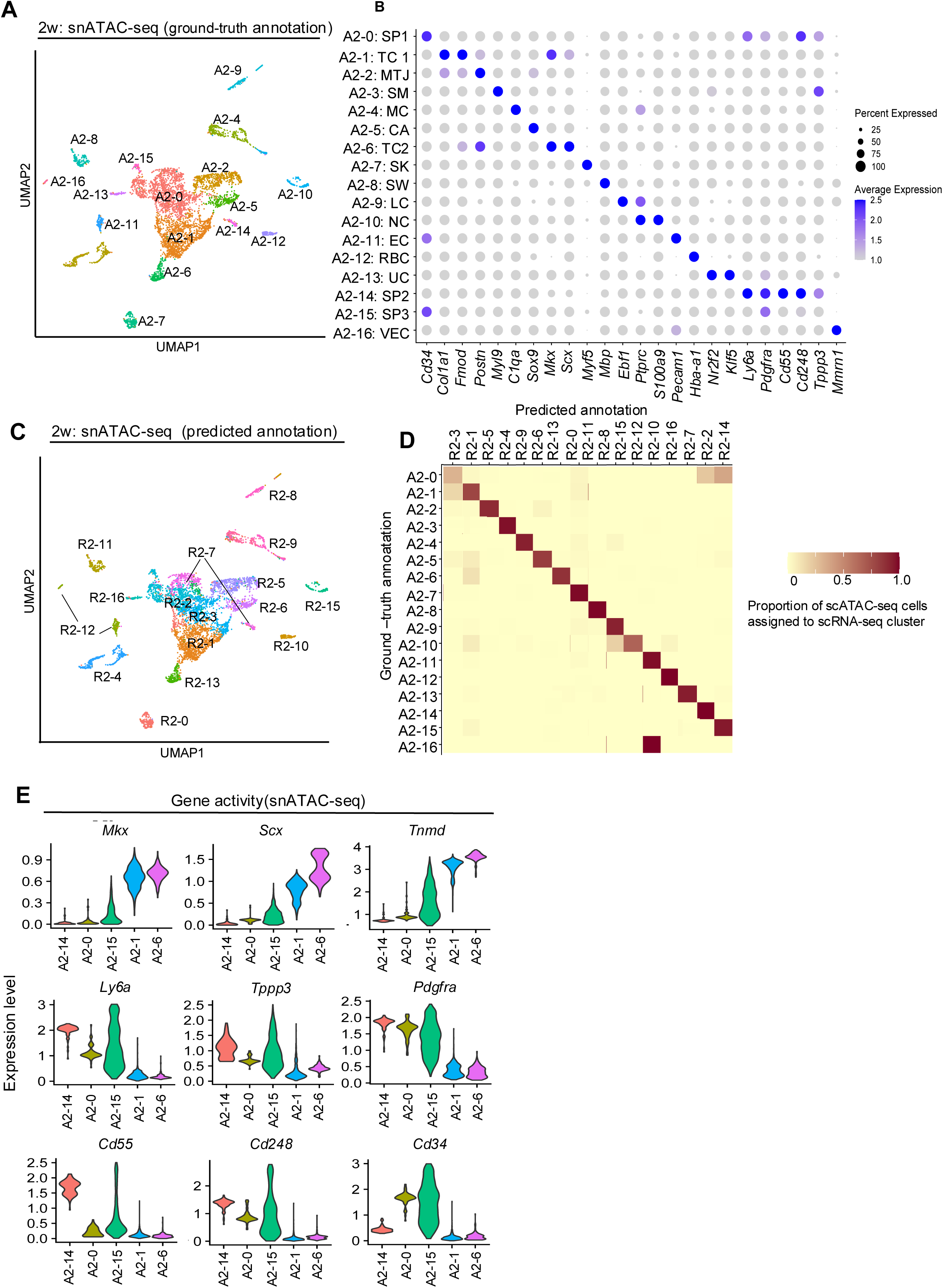
snATAC-seq of tendon cells from a 2-week-old mouse and the validation of Cd55 and Cd248 as candidate markers of TSPC. (A) UMAP snATAC-seq clustering of cells derived from the Achilles tendon of a 2-week-old mouse. Annotation was based on each gene activity (ground-truth annotation). SP1, tendon stem/progenitor cell_1; TC1, tenocyte_1; MTJ, myotendinous junction cell; SM, smooth muscle cell; MC, macrophage; CA, cartilage; TC2, tenocyte_2; SK, skeletal muscle cell; SW, Schwann cell; LC, lymphocyte; NC, neutrophil; EC, endothelial cell; RBC, red blood cell; UC, unspecified cell; SP2, tendon stem/progenitor cell_2; SP3, tendon stem/progenitor cell_3; VEC, vascular endothelial cell. (B) Dot plot of average gene activity of the indicated genes in each snATAC-seq cluster. The size of the dot reflects the percentage of cells in the cluster that express each gene. (C) UMAP visualization and predicted annotation of 2-week snATAC-seq after integration and label transfer of 2-week snRNA-seq data. (D) Identification of matching cell clusters between the 2-week snRNA-and 2-week snATAC-seq data from visualized as heatmap. The heatmap shows the proportions of cells from each snATAC-seq cluster across all sample conditions assigned to each snRNA-seq cluster as part of the label-transfer process. (E) Violin plot of tenocytes and TSPC-related gene expression in each cluster.

Next, we analyzed the snRNA-seq data. We evaluated 10,314 cells (median 3,167 genes/cell and 41,611 mean reads/cell). After doublet removal, we performed unbiased clustering using Seurat and identified 17 clusters (Supplemental Figure 4A). Clusters expressing tendon-related genes *Mkx* and *Scx* were clusters 1 and 13, defined as R2-1: TC_1 and R2-13: TC_2, respectively. Clusters 2, 3, and 7 were identified as expressing stem/progenitor-related genes and defined as R2-2: SP_1, R2-3: SP_2, and R2-7: SP_3, respectively. To validate the consistency of our approaches, we conducted a comprehensive comparison between the cell clusters identified in our scRNA-seq analysis (which included both 2-week and 6-week samples) and our snRNA-seq analysis. We confirmed that all clusters identified in the scRNA-seq data could be similarly annotated in the snRNA-seq data, demonstrating strong concordance between these complementary approaches (Supplemental Figure 5). This validation supports the reliability of our cell type identification across different single-cell methodologies. Interestingly, R2-5: The MTJ cluster expressed not only *Tnc* but also *Sox9*, consistent with previous reports (Nagakura et al., 2020). Using the Signac package’s “FindTransferAnchors” function, we calculated predicted IDs for snATAC-seq clusters based on snRNA-seq annotations (predicted annotation) (Fig. 2C). Evaluation of the two annotation methods for snATAC-seq revealed that both methods allowed annotation of major cell types, and correlation between the two was maintained. Thus, the validity of predicted annotations based on snRNA-seq was confirmed (Figure 2D).

R2-7: SP3, considered the most undifferentiated stem/progenitor fraction in snRNA-seq, could be divided into A2-0: SP1 and A2-14: SP2 in snATAC-seq.

Comparing gene activity of tendon-related and stem/progenitor genes in each cluster of the ground truth annotation, A2-14: SP2 showed high expression of *Ly6a*, *Pdgfra*, and *Tppp3*, and inverse correlation with *Mkx*, *Scx*, and *Tnmd*. Furthermore, the newly identified surface antigens *Cd55* and *Cd248* also showed high expression in cluster 14. In contrast, *Cd34* did not show a high expression pattern in A2-14: SP2. These results suggest that A2-14: SP2 is the most immature cluster, with *Cd55* and *Cd248* showing characteristic expression (Fig. 2E). Subcluster analysis of R7, considered the most immature fraction in snRNA-seq, revealed two clusters based on *Cd55*/*Cd248* and *Cd34* expression, similar to the snATAC-seq analysis. The former (R7-1) showed high expression of *Ly6a*, *Tppp3*, and *Pdgfra* (Fig. 2F). Trajectory analysis using R2-7 and A2-14 as roots in snRNA-seq and snATAC-seq, respectively, showed increased expression of tendon-related genes *Mkx* and *Scx* along the trajectory, while expression of *Tppp3*, *Cd55*, and *Cd248* decreased (Figure 3A). Evaluation of peaks in each cluster based on gene expression levels in snRNA-seq revealed multiple peaks with heights correlated to gene expression for *Mkx* and *Scx*. Similar evaluation of *Cd55* and *Cd248* showed multiple peaks upstream of the TSS for *Cd248* (Fig. 3B). These results indicate correlation between snRNA-seq and snATAC-seq, and consistent with scRNA-seq results, suggest that A2-14 represents the most undifferentiated TSPCs based on *Cd55* and *Cd248* expression.

**Figure 3.**
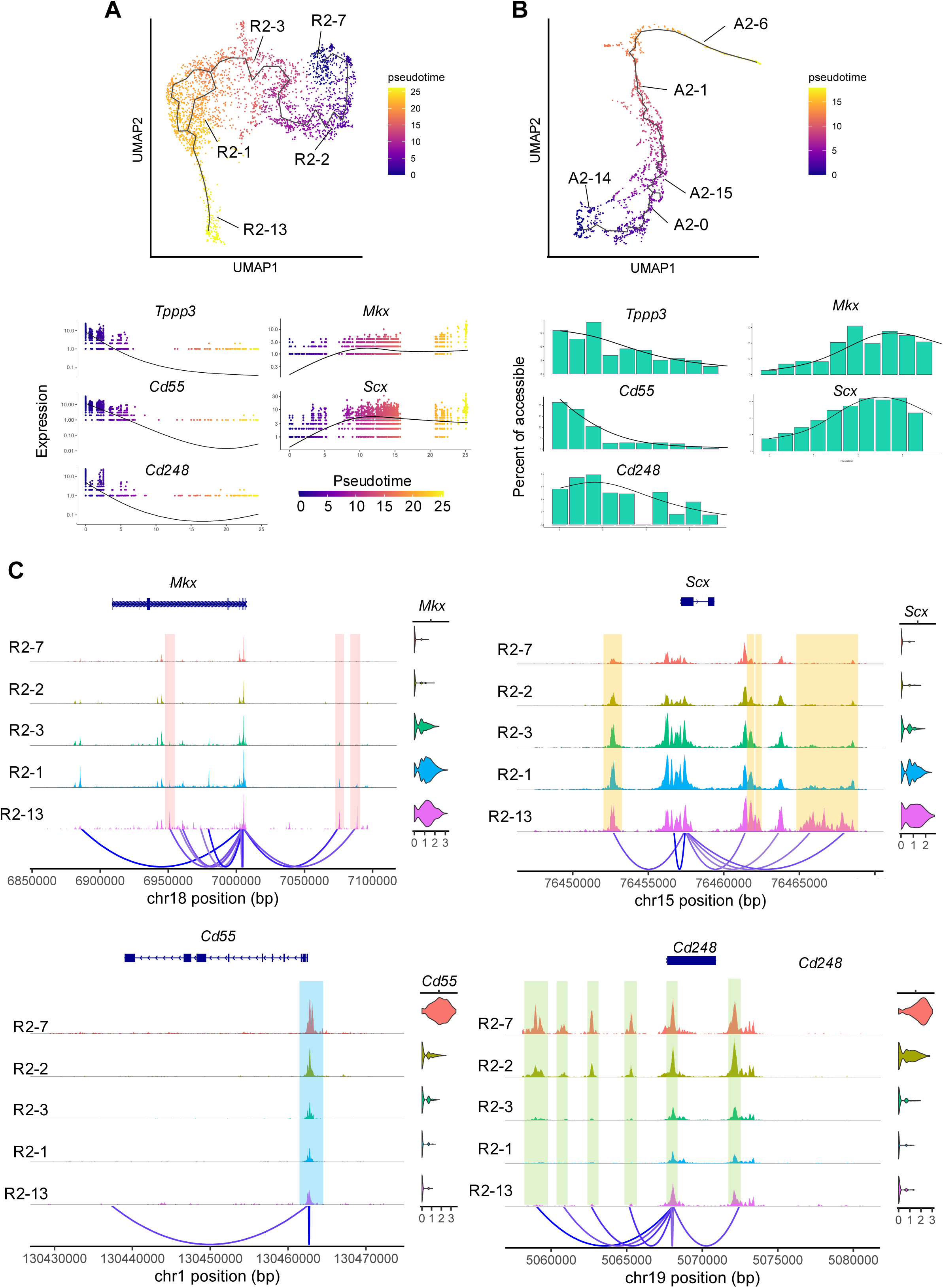
Trajectory analysis and peak visualization of snRNA-seq and snATAC-seq data for the tendon and tendon stem/progenitor cell-related cluster. (A) UMAP representation of snRNA-seq differentiation trajectory of tenocytes and TSPC lineage and pseudotime-dependent gene expression changes of *Tppp3*, *Cd55*, *Cd248*, *Mkx,* and *Scx,* as inferred using Monocle3. (B) UMAP representation of snATAC-seq differentiation trajectory of tenocytes and the TSPC lineage and pseudotime-dependent gene expression changes, as inferred using Cicero. (C) Coverage plots of *Mkx*, *Scx*, *Cd55,* and *Cd248*. Selected peaks that differ across each cluster are highlighted.

### Comparison of Single Nucleus RNA+ATAC Analysis between 2-week and 6-week Mouse Achilles Tendon Cells

To confirm the presence of the identified *Cd55*/*Cd248* fraction at 6 weeks, we performed snRNA-seq and snATAC-seq on 6-week-old mouse Achilles tendon cells. Using the 2-week snATAC-seq data as a reference, we annotated the 6-week snATAC-seq clusters. Clusters A2-14 and A2-0 corresponded to clusters A6-12 and A6-0, respectively (Supplemental Figure 6). We also annotated the 6-week snATAC-seq clusters using 2-week snRNA-seq with the same results (Supplemental Figure 7). Gene activity evaluation revealed that, similar to the 2-week data, the 6-week A6-12 and A6-0 clusters showed *Cd55*/*Cd248* high and *Cd34* dim, and Cd55/*Cd248* dim and *Cd34* high patterns, respectively. Furthermore, annotation of the 6-week snRNA-seq data using the 2-week snRNA-seq as a reference revealed clusters with expression patterns similar to those observed in snATAC-seq. In both cases, clusters with high *Cd55* and *Cd248* expression showed high gene activity (snATAC-seq) and gene expression (snRNA-seq) of *Tppp3*, *Pdgfra*, and *Ly6a*. These results confirm that the *Cd55* and *Cd248* high clusters identified in the 2-week snRNA-seq+snATAC-seq analysis are similarly detected at 6 weeks.

We also compared the snATAC-seq data between 2 and 6 weeks. No significant differences were observed in genomic annotations between the two time points (Supplemental Figure 8A). After merging the datasets (Supplemental Figure 8B), we compared gene activity of tendon/stem-related genes to evaluate changes in gene activity. We found that the activity of *Tppp3*, *Pdgfra*, *Ly6a*, *Cd55*, *Cd248*, and *Cd34* all decreased at 6 weeks (Supplemental Fig 8C). This was consistent with the decreased gene expression observed when comparing 2-week and 6-week data.

### Estimation of Transcription Factor Activity in 2-week Mouse Achilles Tendon Cells

To estimate transcription factor activity in each cluster, we used the Single-Cell Regulatory Network Inference and Clustering (SCENIC) package (Lake et al., 2018) to calculate gene regulatory network activity from scRNA-seq gene expression data. SCENIC constructs gene expression networks centered on transcription factors and infers transcription factor activity in each cluster. We summarized the predicted transcription factor activities in tendon and stem/progenitor-related clusters R2-7, R2-2, R2-3, R2-1, and R2-13 identified by snRNA-seq (Figure 4).

**Figure 4.**
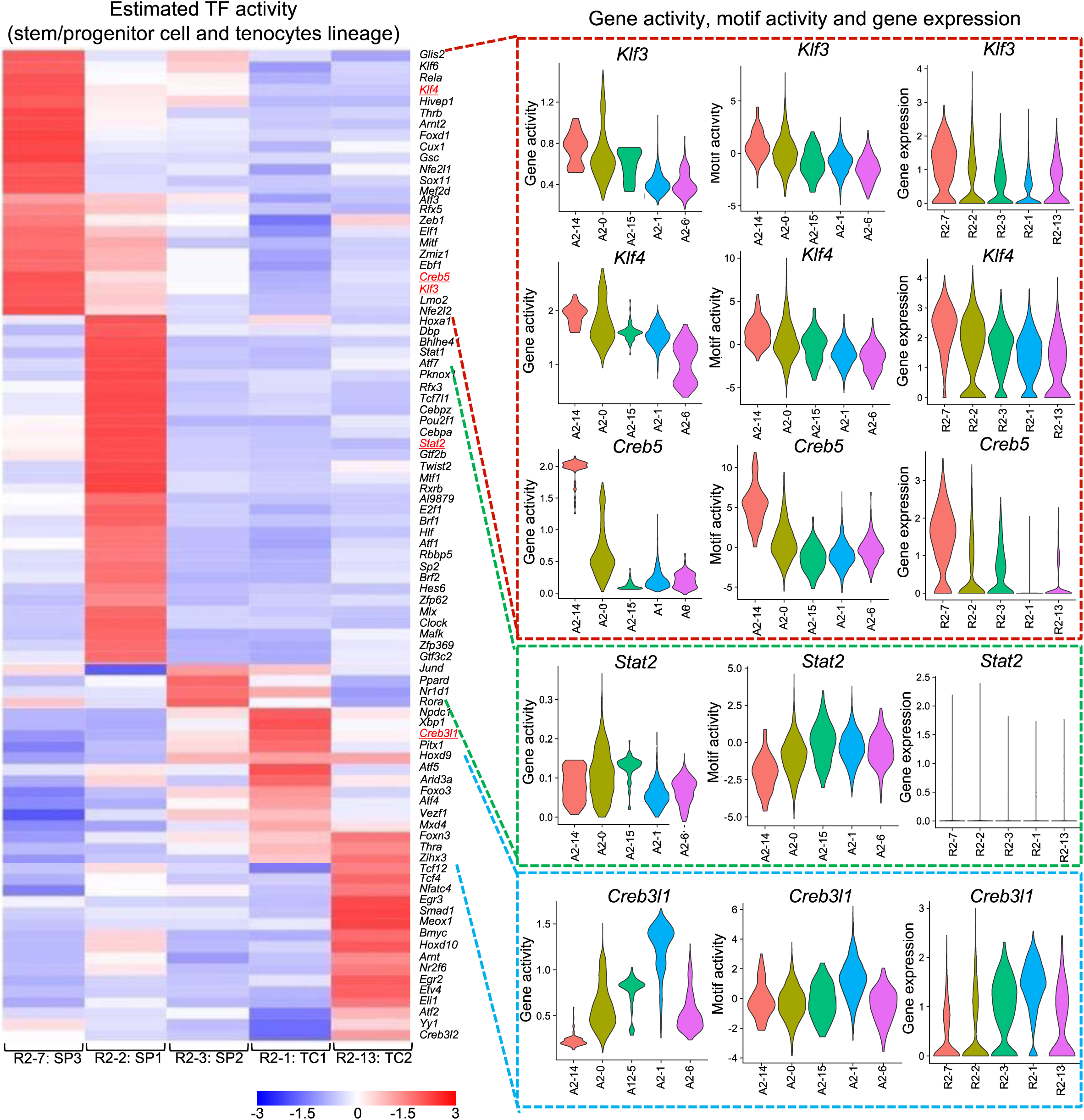
Transcription factor landscapes of 2-week mouse Achilles tendons. SCENIC analysis of transcription factor activity based on 2-week snRNA-seq data for tenocytes and the TSPC lineage (Left). Validation was performed based on the gene activity and motif activity of 2-week snATAC-seq and gene expression of 2-week snRNA-seq (right).

Next, we evaluated the gene activity of these identified transcription factors in each snATAC-seq cluster, along with motif activity calculated by chromVAR (Schep et al., 2017). We also assessed the expression levels of transcription factors in each snRNA-seq cluster. In the most immature fractions, represented by cluster A2-14 in snATAC-seq and cluster R2-7 in snRNA-seq, *KLF transcription factor 3* (*Klf3*), *KLF transcription factor 4* (*Klf4*), and *cAMP responsive element binding protein 5* (*Creb5*) showed consistent behavior in gene activity, motif activity, and gene expression. Additionally, *Signal transducer and activator of transcription 2* (*Stat2*) and cAMP responsive element binding protein 3 like 1 (*Creb3l1*) showed high values in A2-0/A2-15 and A2-1, respectively. *Stat2* gene expression was not observed, likely due to false positives resulting from low expression levels. The gene activity and motif activity of each transcription factor estimated by snATAC-seq correlated with the transcription factor activity calculated by SCENIC. Therefore, simultaneous analysis of snRNA-seq and snATAC-seq could be used to more accurately evaluate the function of transcription factors in each cluster.

Furthermore, SCENIC can predict candidate transcription factors regulating each gene (Table 1). *Cd55* and *Cd248* were predicted to be regulated by *Klf3* and *Klf4*, while *Mkx* and *Scx* were predicted to be under the control of *Creb3l1*. These predictions were consistent with the data calculated from gene activity and motif activity in snATAC-seq.

**Table1:**
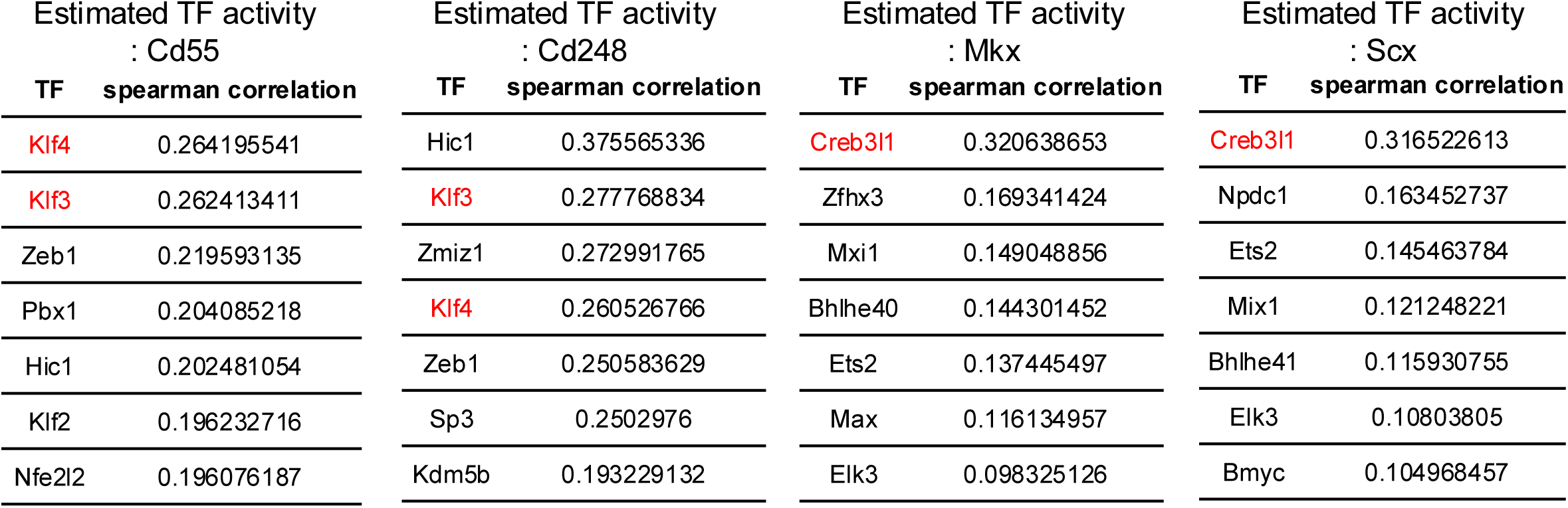
Estimated TF activity in each gene.

### *In vitro* Evaluation of CD55 and CD248

Immunostaining evaluation of CD55 and CD248, the identified candidate stem/progenitor markers, revealed expression in the peritenon (Figure 5C, 5A). This was consistent with previous reports of localized expression of *Tppp3*/*Pdgfra*-positive cells in the peritenon and our analysis showing co-expression of *Tppp3*/*Pdgfra* and *Cd55*/*Cd248*. The results support the possibility that SP clusters reflect peritenon. Given that *Cd55* and *Cd248* expression appeared to reflect *Tppp3*, *Pdgfra*, and *Ly6a* expression more sensitively than *Cd34*, we extracted tendon cells from 2-week-old mice and sorted them by FACS to determine the biological phenotype of Cd55 and Cd248 positive cells (Figure 5B).

**Figure 5.**
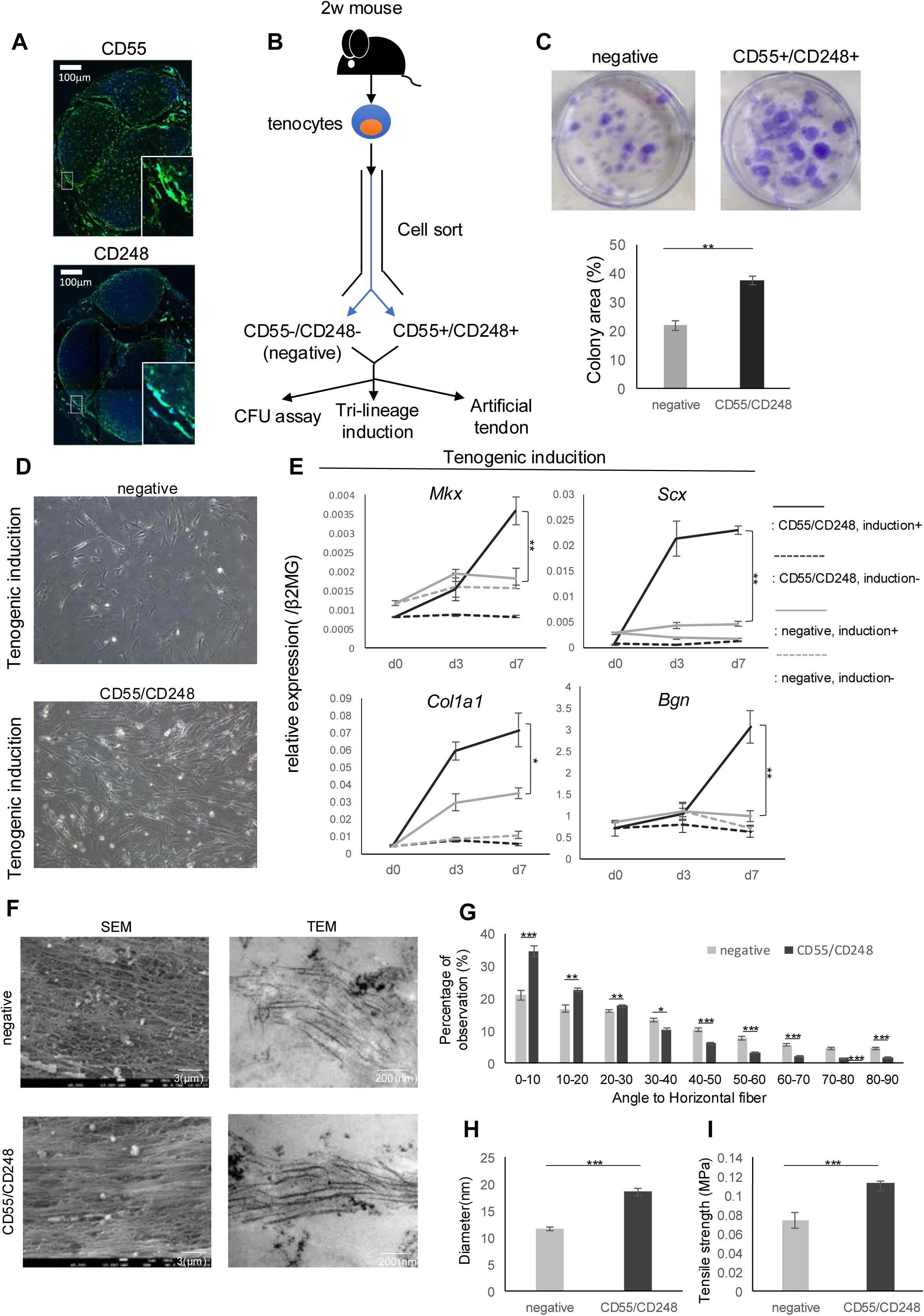
*In vitro* analysis of CD55+/CD248+ TSPCs. (A) Immunohistochemical image of 10-week mouse Achilles tendons. Scale bars show 100μm. CD55 and CD248, green; Hoechst 33342, blue. (B) Schema of the *in vitro* assessment of the capacity of CD55+/CD248+ TSPCs as the differentiation toward tenocytes. (C) Colony-forming efficiency of CD55+/CD248+ and CD55-/CD248-(negative) TSPCs. Colonies were stained with crystal violet (n = 6). CD55+/CD248+ TSPC exhibited higher clonogenic capacity. Data are presented as means ± SEM. \*\**p* < 0.01. (D) Morphological changes of CD55+/CD248+ and negative TSPCs after tenogenic induction. (E) Quantitative PCR of tendon-related gene expression in CD55+/CD248+ and negative TSPCs after tenogenic induction (n = 3). Data are presented as means ± SD. \*\**p* < 0.01, \**p* < 0.05. (F) SEM and TEM imaging of artificial tendons derived from CD55+/CD248+ and negative TSPCs. Data are presented as means ± SD. \*\**p* < 0.01, \**p* < 0.05. (G) Proportions of fiber alignment for each artificial tendon (n = 4). Data are presented as means ± SEM. \*\*\**p* < 0.005, \*\**p* < 0.01, \**p* < 0.05. (H) Diameter of collagen fiber in each artificial tendon based on TEM imaging (n = 4). Data are presented as means ± SD. \*\*\**p* < 0.005. (I) Tensile strength (MPa) of each artificial tendon (n = 5). Data are presented as means ± SEM. \*\*\**p* < 0.005.

Gene expression analysis of sorted cells showed that CD55/CD248 positive cells, compared to negative cells, had lower expression of *Mkx*, *Scx*, *Col1a1*, and *Creb3l1*, but higher expression of *Ly6a*, *Tppp3*, *Pdgfra*, *Creb5*, and *Klf3*, consistent with snRNA-seq analysis (Supplemental Figure 9). To assess stem/progenitor capacity, we performed colony formation assays. CD55/CD248 sorted cells showed significantly increased colony formation compared to CD55/CD248 negative cells (control) (Figure 5C).

We then evaluated the differentiation tendencies of these cells towards tendon, cartilage, and bone. Tendon differentiation resulted in clusters of spindle-shaped cells from CD55/CD248 positive cells, suggesting tenogenic differentiation (Figure 5D). Gene expression analysis revealed that while CD55/CD248 positive cells initially had lower *Mkx* and *Scx* expression than negative cells, this pattern reversed after differentiation (Figure 5E). Increased expression of other tendon-related genes such as *Col1a1* and *Bgn* was also observed. In contrast, no significant increases in *Sox9* and *Collagen type II alpha 1 chain* (*Col2a1*) (chondrogenic) or *RUNX family transcription factor 2* (*Runx2*), and *Alkaline phosphatase, biomineralization associated* (*Alpl*) (osteogenic) expression were observed compared to negative cells during cartilage and bone differentiation assays (Supplemental Figure 10). Cells negative for CD55/CD248 could be mixed cell populations, including hematopoietic lineages, cells from tendon mid substance, immune cells, and/or endothelial cells. However, given that CD55/CD248 negative cells have increased expression of genes characteristic of tendons such as *Mkx*, *Scx* and *Col1a1*, we could speculate that this popularity might reflect terminally differentiated tenocytes and CD55/CD248 positive cells possess tenogenic differentiation capacity.

To further evaluate the tenogenic potential of CD55/CD248 positive cells, we created tendon-like tissue (bio-tendon) using our previously reported 3D stretch stimulation culture system. SEM and TEM analysis to assess collagen fiber density and thickness showed that bio-tendons derived from CD55/CD248 positive cells had an increased proportion of collagen fibers parallel to the stretch direction and increased collagen fiber diameter. TEM also revealed characteristic banding patterns and triple helix structures in these bio-tendons, indicating mature collagen organization (Wieczorek et al., 2015) (Figure 5F-H).

Stretch tests to evaluate the mechanical capacity of the bio-tendons showed high tensile strength (Figure 5I).

These results demonstrate that CD55/CD248 positive cells have a tendency to differentiate into tendon cells.

## Discussion

Tendons are known for their low cellular content, making complete functional recovery after injury challenging and increasing the risk of re-rupture. As a result, cell therapy has garnered attention as a novel treatment strategy, distinct from current conservative and surgical approaches. However, this requires a thorough understanding of TSPCs. Research on TSPCs has been limited due to the lack of identified characteristic surface antigens.

To our knowledge, no multi-omics analysis comparing juvenile and mature mouse tendon cells has been conducted. In this study, we performed single-cell RNA-seq along with snRNA-seq+snATAC-seq on nuclei isolated from 2-week and 6-week mouse Achilles tendons. While comparisons using older mice (e.g., 12 weeks or more) were initially considered, they were excluded due to extremely low cell yield and viability, making 2-and 6-week-old mice more suitable for this analysis. This approach allowed us to identify candidate surface antigens for TSPCs and elucidate the dynamism of transcription factors involved in post-developmental tendon growth. As a result, we discovered a novel combination of *Cd55* and *Cd248* as surface antigens showing characteristic expression in the TSPC fraction.

While scRNA-seq analysis also showed *Cd34* expression characteristic of the TSPC fraction, consistent with previous reports, snRNA-seq + snATAC-seq analysis revealed that the *Cd34* high-expression cluster could be further classified into two clusters based on *Cd55* and *Cd248* expression patterns. The cluster showing high expression of both *Cd55* and *Cd248* also exhibited high expression of *Tppp3*, *Pdgfra*, and *Ly6a*, suggesting that isolating *Cd55* and *Cd248* positive fractions may more sensitively capture immature populations compared to *Cd34*.

snRNA-seq+snATAC-seq analysis provided simultaneous information on gene expression levels and open chromatin regions for each cluster. Notably, *Cd248* showed several peaks upstream of the TSS in high-expression clusters, likely reflecting its expression regulation mechanism. We also identified several peaks that increased proportionally with expression for *Mkx* and *Scx*. Given the many unknowns in *Mkx* and *Scx* expression regulation mechanisms (Guerquin et al., 2013; Otabe et al., 2015), we used the SCENIC package to predict upstream transcription factors based on expressed genes in each snRNA-seq cluster. Combined with gene activity and motif activity data from snATAC-seq, Creb3l1 was identified as a transcription factor regulating *Mkx* and *Scx* expression. *Creb3l1* has been reported to increase in expression throughout development (Liu et al., 2015).

*Klf3*, *Klf4*, and *Creb5* were identified as characteristic transcription factors in the TSPC fraction. Recent reports have highlighted the role of the *Klf* family in limb development (Kult et al., 2021). Given that tendons are components of the developing limb, these transcription factors may have roles in tendon biology. *Creb5* has been reported to show increased expression from E11 to E13 (Liu et al., 2015), suggesting a role in early developmental stages. Furthermore, *Creb5* has been reported to regulate Proteoglycan 4 (*Prg4*) expression (Zhang et al., 2021). Further investigation into the functions of these transcription factors in TSPCs is necessary.

CD55 was identified by Hoffmann et al. in 1969 as a surface antigen functioning as a complement inhibitory factor on erythrocytes. CD55 is known to be expressed from early developmental stages and characterizes the initial differentiation stage of hematopoietic stem cells (Guo et al., 2013). It is also expressed in MSCs and has been reported as an early progenitor marker for mouse mammary epithelial cells (Pal et al., 2017). CD248 was identified in 1992 as an antigen for the FB5 antibody reacting with vascular wall cells (Rettig et al., 1992). CD248 is a transmembrane glycoprotein expressed in pericytes and fibroblasts during developmental stages. Regarding their relationship, CD55 and CD248 have been reported to be expressed in stromal cells during the early stages of arthritis (Choi et al., 2017), but their detailed mechanisms in tendons remain unclear.

Previously suggested TSPC surface antigen candidates such as *Cd73*, *Cd90*, and *Cd105* showed poor correlation with the expression of genes like *Tppp3*. Moreover, the initial report on TSPCs (Bi et al., 2007) described TSPCs as *Scx*+*Cd34*-. The high expression of CD55 and CD248 in the peritenon, similar to *Tppp3* (Harvey et al., 2019; Staverosky et al., 2019), suggests the possibility of cells with tenogenic differentiation potential exhibiting *Scx*+*Cd34*-expression patterns within the tendon. Spatial information is important to investigate further. Future studies using lineage tracing experiments with mice labeled for CD55 and CD248 or mouse model of selective ablation of CD55 and CD248 are necessary to analyze the developmental functions of CD55 and CD248 positive cells and their roles in injury healing in more detail.

Clinically, to our knowledge, few studies have sorted TSPCs based on surface antigens and examined their *in vitro* tendon tissue-generating ability. In this study, we found that cells sorted for CD55/CD248 showed higher clonogenicity compared to CD55-/CD248-cells and demonstrated superior tendon tissue-generating ability in an artificial tendon model. In the future, CD55/CD248 double-positive TSPC cells may prove useful in clinical applications such as *in vitro* artificial tendon creation (Tsutsumi et al., 2022) and cell therapy for tendon injuries (Huang et al., 2021).

## Acknowledgements

We thank all the members of the Department of Systems BioMedicine at Institute of Science Tokyo for their support. We also thank the Research Core at Institute of Science Tokyo for supporting cell sorting.

## Author contributions

Conceptualization, H.T., T.C., and H.A.; Methodology, H.T., T.C., and H.A.; Formal analysis, H.T.; Investigation, H.T., F. Y., T. M., T. K., A. K., K. A., Y. S.; Writing – Original Draft, H.T.; Writing – Review & Editing, T.C., and H.A.; Funding Acquisition, H.A.; Supervision, H.A.

## Data availability

FASTQ data of RNA-Seq and ATAC-seq are deposited in DDBJ under accession number PRJDB18857

---

**Supplemental Figure 1.**
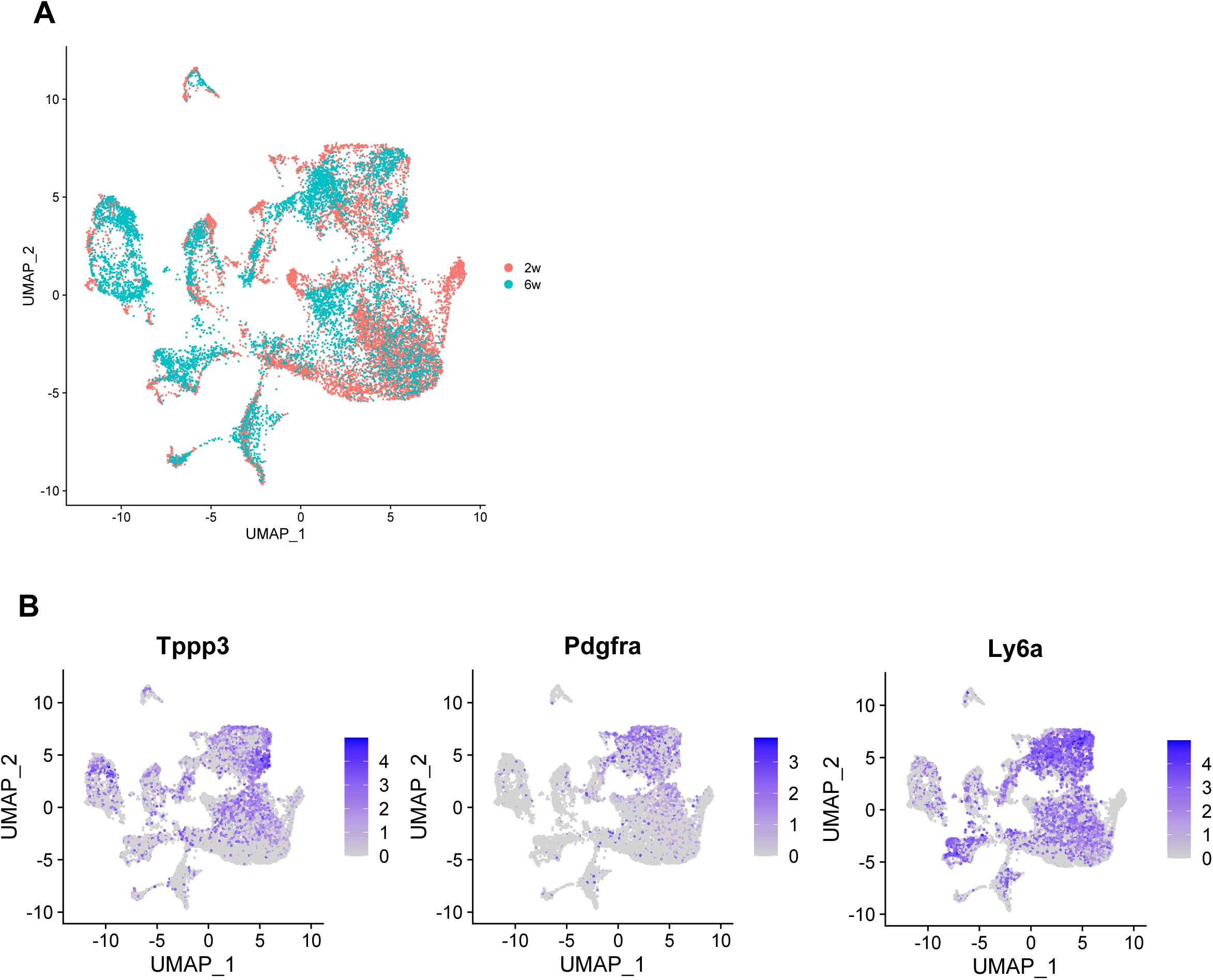
2w-and 6w-scRNA seq results. (A) Integrated uniform manifold approximation and projection (UMAP) scRNA-seq clustering of cells harvested from 2-week-old and 6-week-old mouse Achilles tendons. Clusters are colored according to cluster type. (B) Feature plot of *Tppp3*, *Pdgfra* and *Ly6a* expression.

**Supplemental Figure 2.**
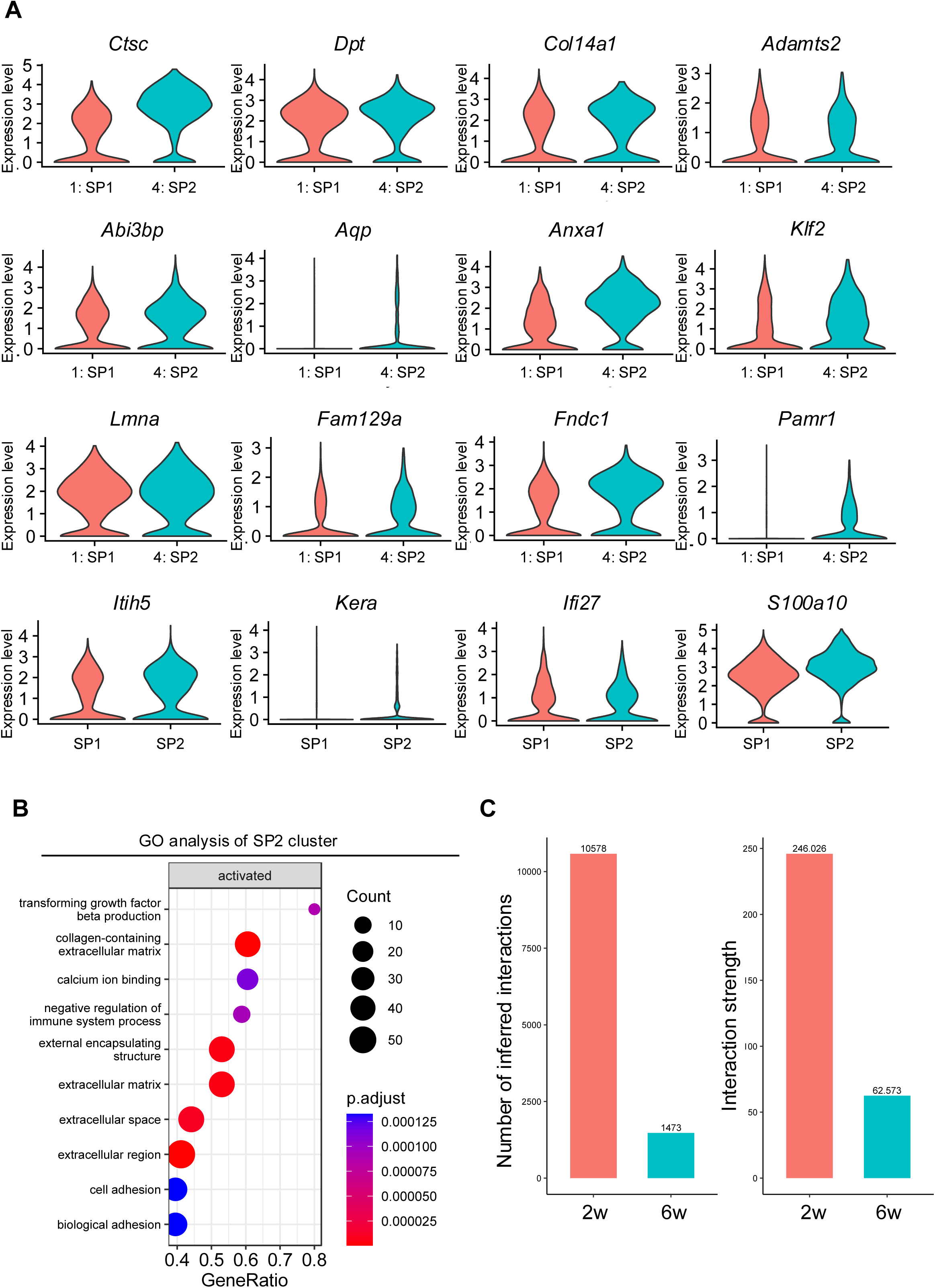
Comparison of 2w-and 6w-scRNA seq results. (A) Violin plot of gene expression enriched in the limb bud (Development (2014) 141, 3683-3696) in each cluster. (B) Gene Ontology (GO) terms associated with genes with upregulated expression in the SP2 cluster. (C) Number of inferred interactions and interaction strength of genes expressed within 2w-and 6w-mouse scRNA-seq data.

**Supplemental Figure 3.**
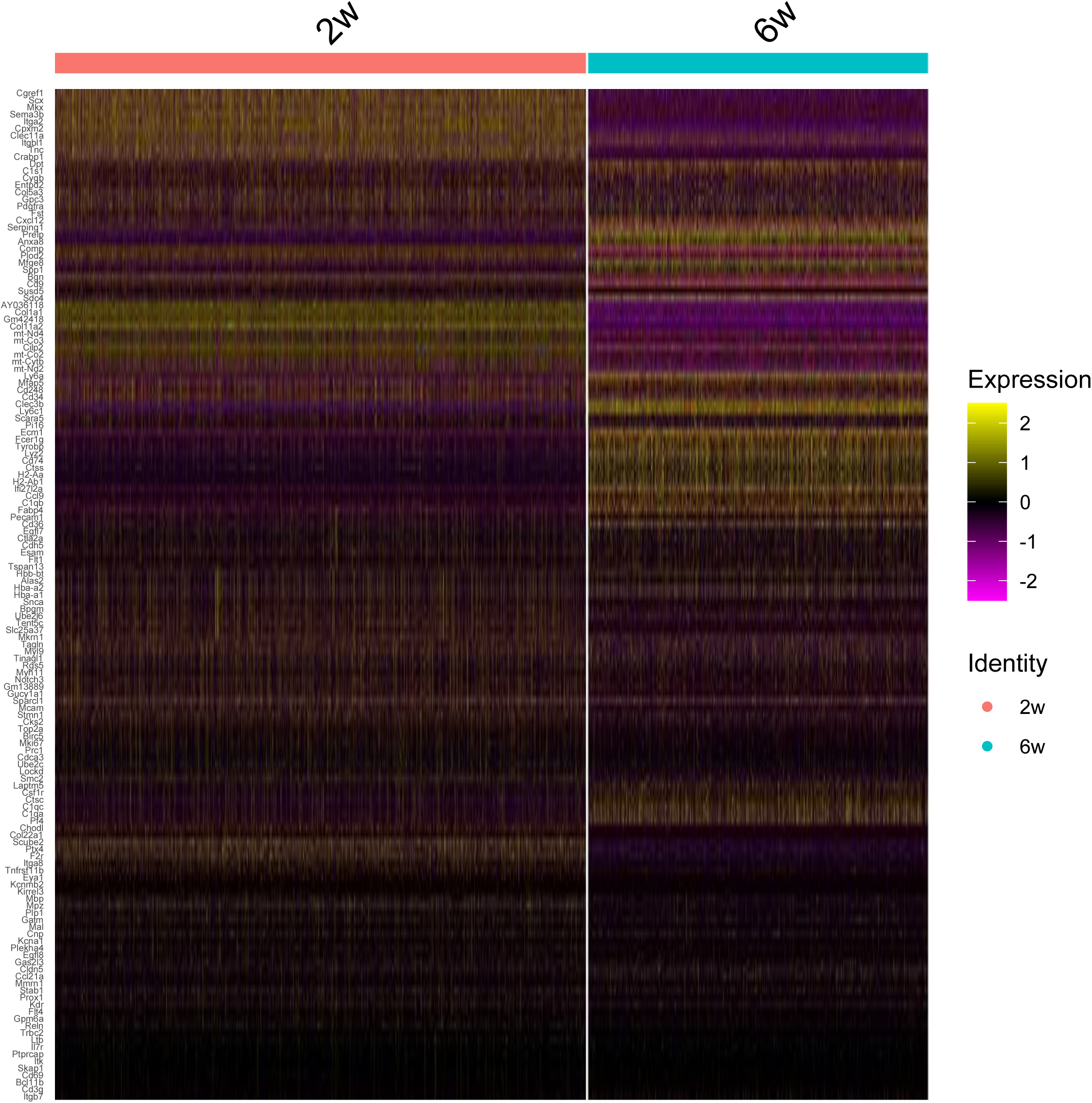
Comparative analysis of 2w-and 6w-scRNA seq results. Heatmap showing differential gene expression between red (2w) and green (6w).

**Supplemental Figure 4.**
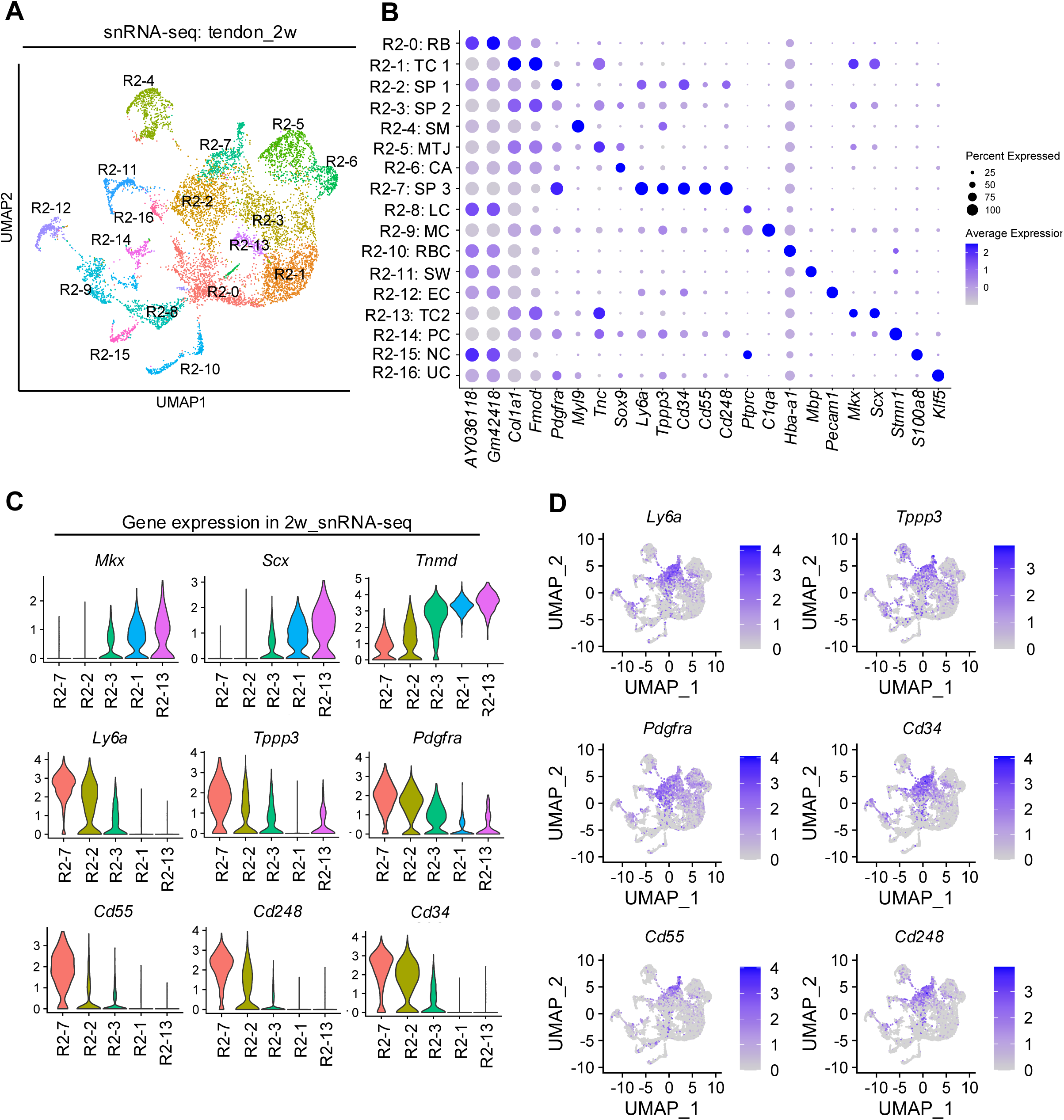
Analysis of 2-week snRNA-seq data. (A) UMAP snRNA-seq clustering of cells harvested from 2-week mouse Achilles tendons. (B) Dot plot of average gene expression levels of the indicated genes in 2-week snRNA-seq clusters. RB, ribosomal RNA; TC1, tenocyte_1; SP1, stem/progenitor cell_1; SP2, stem/progenitor cell_2; SM, smooth muscle cell; MTJ, myotendinous junction cell; CA, cartilage; SP3, stem/progenitor cell_3; LC, lymphocyte; MC, macrophage; RBC, red blood cell; SW, Schwann cell; EC, endothelial cell; TC2, tenocyte_2; PC, proliferating cell; NC, neutrophil; UC, unspecified cell. (C) Violin plot of tenocytes and TSPC-related gene expression in each cluster. (D) Feature plot of TSPC-related gene expression.

**Supplemental Figure 5.**
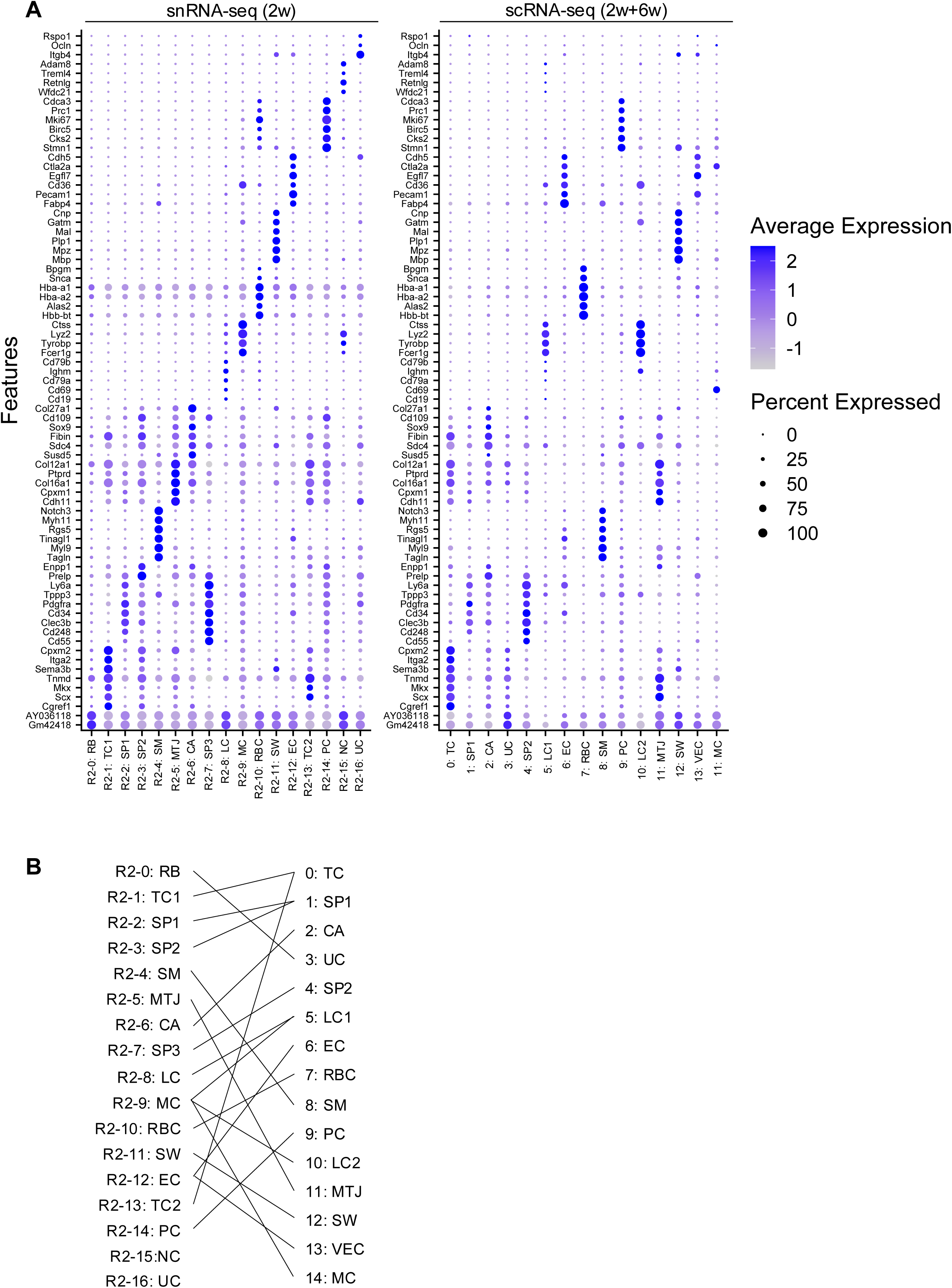
Analysis of 6-week snATAC-seq. (A) UMAP snATAC-seq clustering of cells harvested from 6-week mouse Achilles tendons. Annotation was based on gene activity (ground-truth annotation, left) and the predicted annotation inferred from 2-week snATAC-seq (right). (B) Identification of matching cell clusters between the 2-week and 6-week snATAC-seq data, visualized as a heatmap. (C) Violin plot of tenocytes and TSPC-related gene activity in each cluster.

**Supplemental Figure 6.**
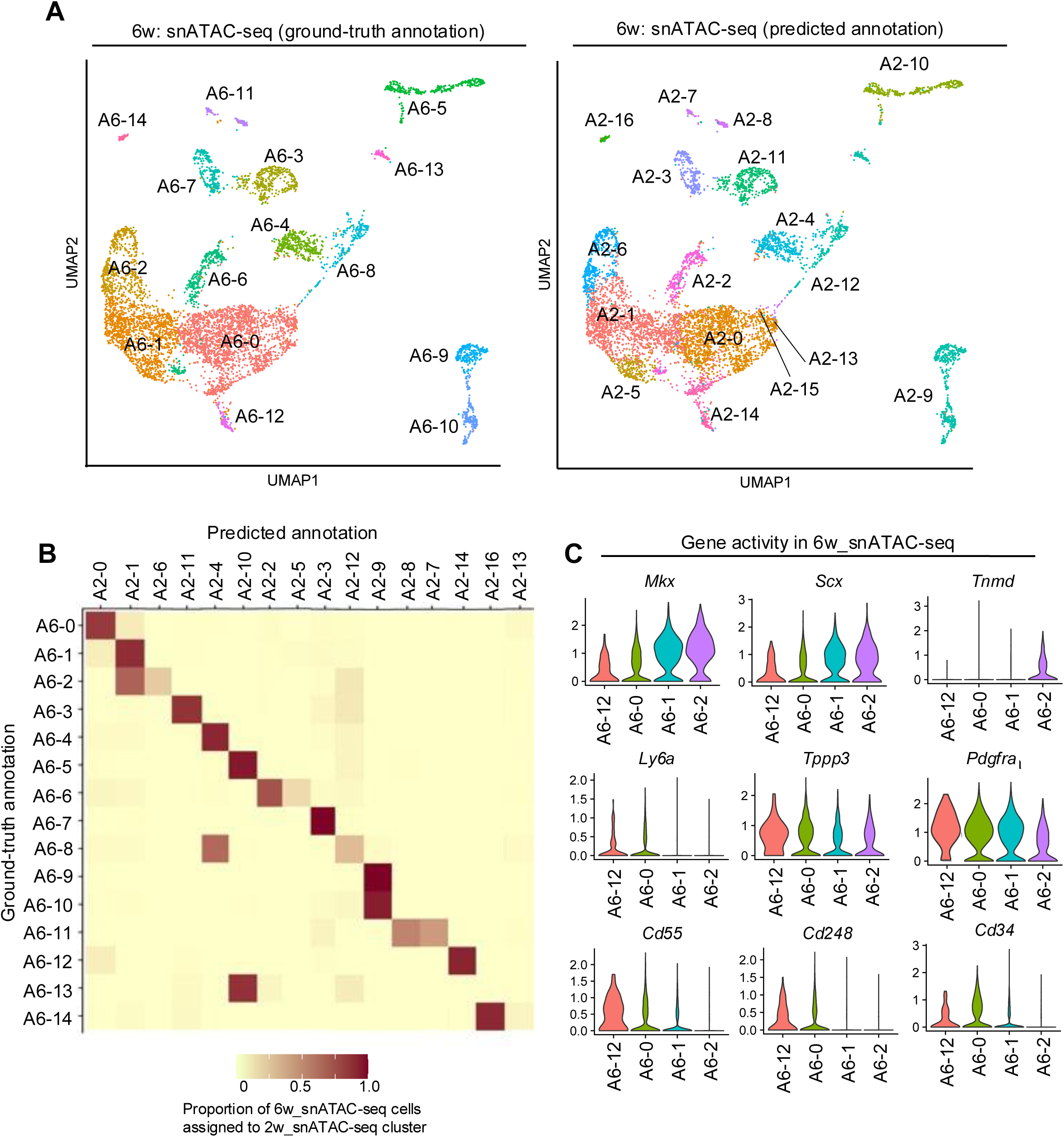
Comparison of snRNA-seq and scRNA-seq. (A) Dot plot of representative DEGs in snRNA-seq (2w) and snRNA-seq (2w+6w). (B) Relevance of annotation in each data.

**Supplemental Figure 7.**
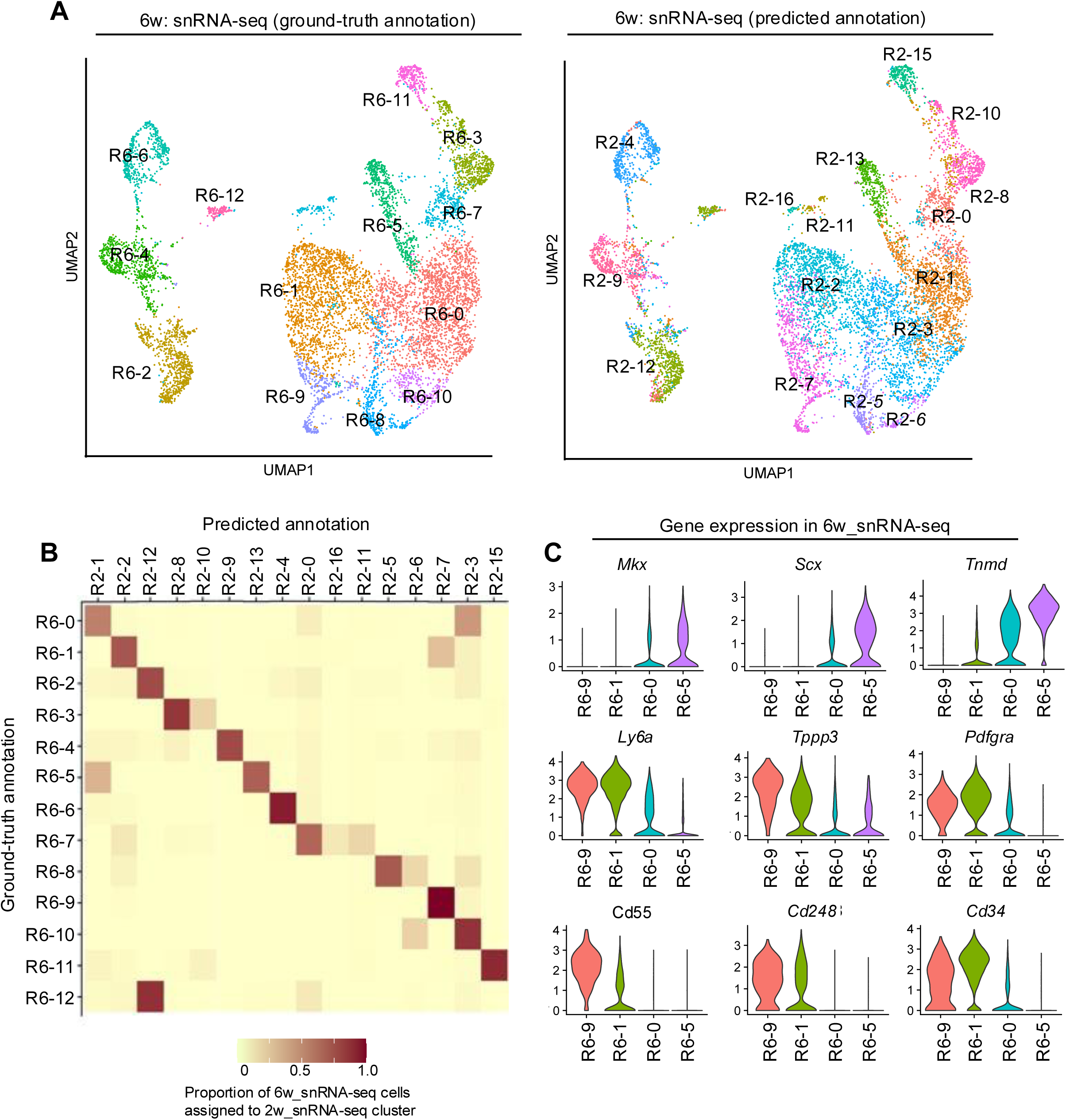
Analysis of 6-week snRNA-seq. (A) UMAP snRNA-seq clustering of cells harvested from 6-week mouse Achilles tendons. Annotation was based on each gene expression (ground-truth annotation, left) and predicted annotation inferred from 2-week snRNA-seq (right). (B) Identification of matching cell clusters between the 2-week and 6-week snRNA-seq data visualized as a heatmap. (C) Violin plot of tenocytes and TSPC-related gene expression in each cluster.

**Supplemental Figure 8.**
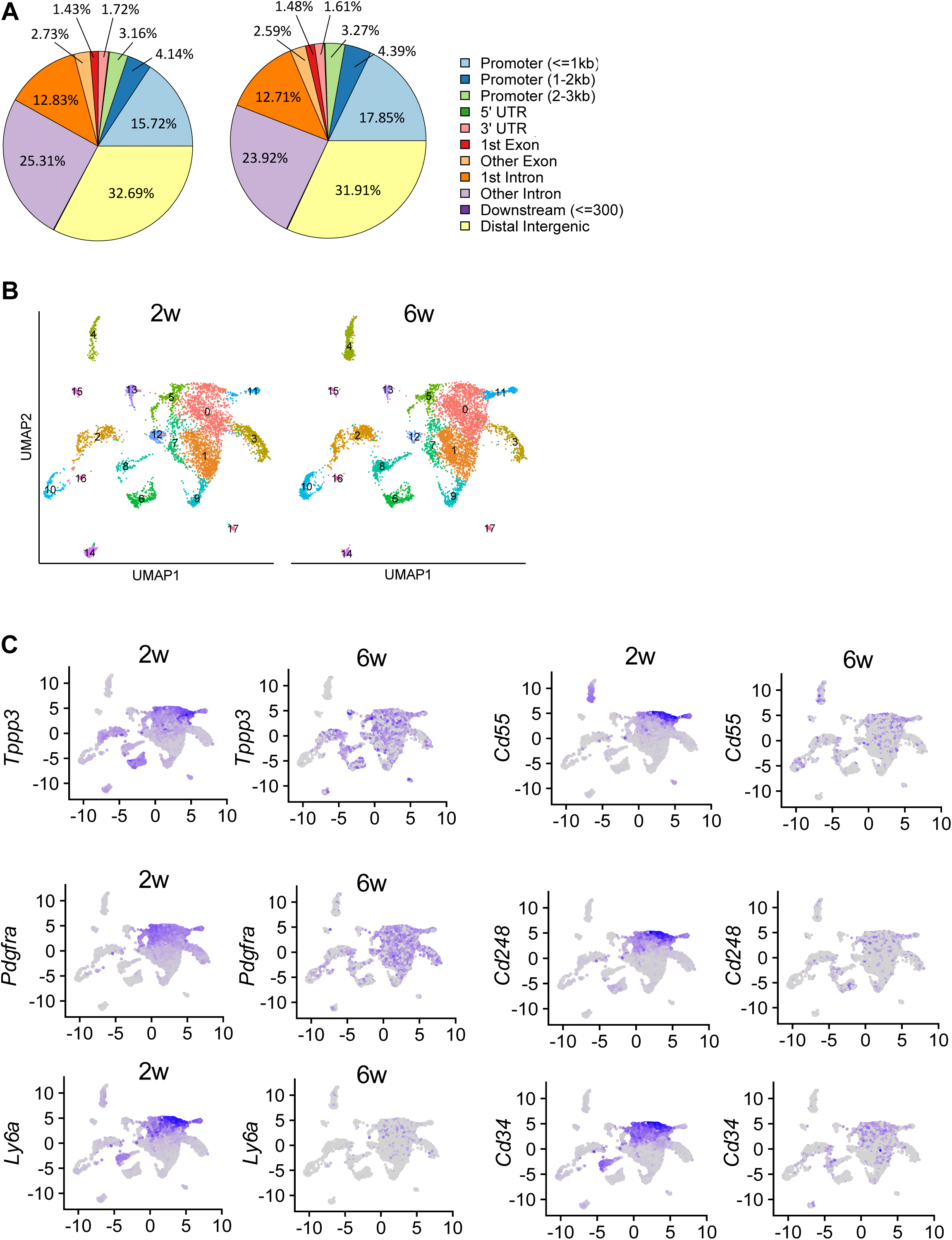
Comparison of 2-week and 6-week snATAC-seq. (A) Circle plot of annotated differentially accessible regions for each data. (B) Integrated UMAP snATAC-seq clustering of cells harvested from 2-week and 6-week mouse Achilles tendons. (C) Feature plot of TSPC-related gene activity in each dataset.

**Supplemental Figure 9.**
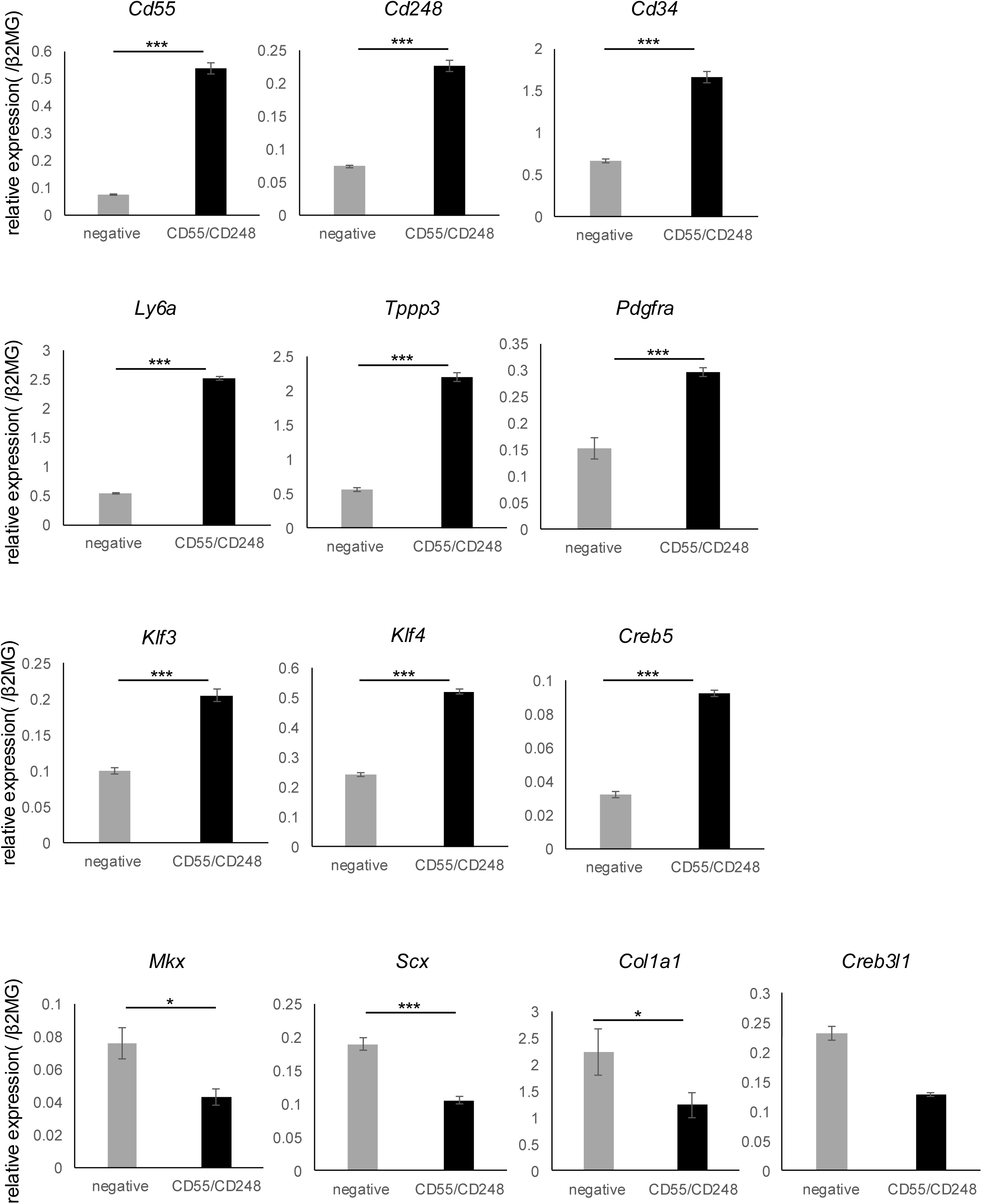
Gene expression changes in CD55+/CD248+ and negative TSPCs. Quantitative PCR of gene expression in CD55+/CD248+ and negative TSPCs (n = 4). Data are presented as means ± SEM. \**p* < 0.05, \*\*\**p* < 0.005.

**Supplemental Figure 10.**
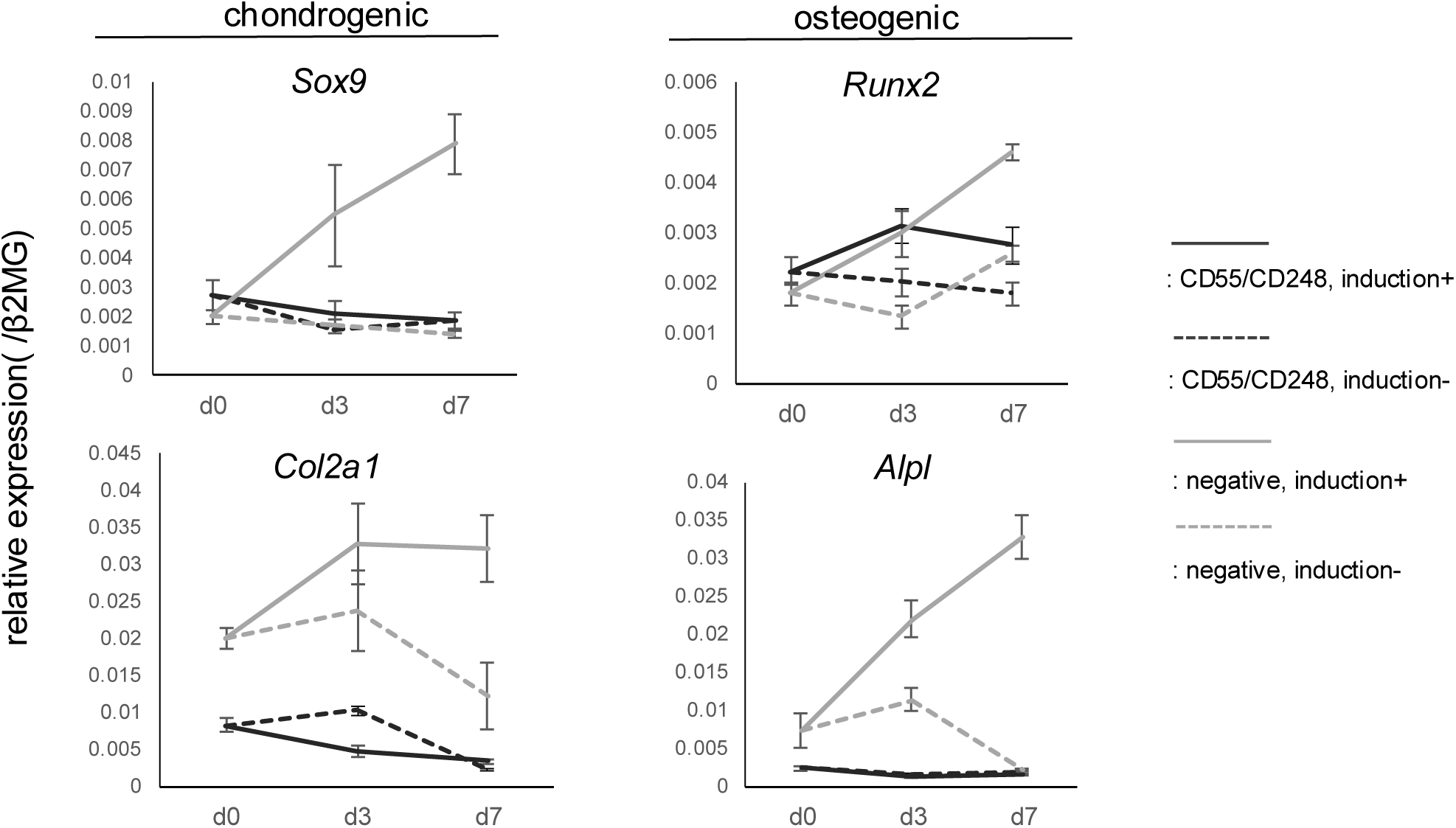
Chondrogenic and osteogenic induction of CD55+/CD248+ and negative TSPCs. Quantitative PCR of tendon-related genes in CD55+/CD248+ and negative TSPCs after chondrogenic and osteogenic induction (n = 4). Data are presented as means ± SEM. \*\**p* < 0.01, \**p* < 0.05.

